# Endometriotic lesions exhibit distinct metabolic signature compared to paired eutopic endometrium at the single-cell level

**DOI:** 10.1101/2024.02.27.580606

**Authors:** Meruert Sarsenova, Ankita Lawarde, Amruta D. S. Pathare, Merli Saare, Vijayachitra Modhukur, Pille Soplepmann, Anton Terasmaa, Tuuli Käämbre, Kristina Gemzell-Danielsson, Parameswaran Grace Luther Lalitkumar, Andres Salumets, Maire Peters

## Abstract

Current therapeutics of endometriosis are limited to hormonal action on endometriotic lesions to disrupt their growth. Based on the recent findings of the high utilization of glycolysis over oxidative metabolism (Warburg-like effect) in endometriotic lesions, a new strategy of nonhormonal management by addressing cellular metabolism has been proposed. However, it remains unclear which cell types are metabolically altered and contribute to endometriotic lesion growth for targeting them with metabolic drugs. Using single-cell RNA-sequencing, we investigated the activity of twelve metabolic pathways and genes involved in steroidogenesis in paired samples of eutopic endometrium (EuE) and peritoneal lesions (ectopic endometrium, EcE) from women with confirmed endometriosis. We detected nine major cell clusters in both EuE and EcE. The metabolic pathways were differentially regulated in perivascular, stromal and to a lesser extent in endothelial cell clusters, with the highest changes in AMP-activated protein kinase signaling, Hypoxia-Inducible Factor-1 signaling, glutathione metabolism, oxidative phosphorylation, and glycolysis/gluconeogenesis. We identified a transcriptomic co-activation of glycolysis and oxidative metabolism in perivascular and stromal cells of EcE compared with EuE, suggesting that metabolic reprogramming may play a critical role in maintaining cell growth and survival of endometriotic lesions. Additionally, progesterone receptor was significantly downregulated in perivascular and endothelial cells of EcE. The expression of estrogen receptor 1 was significantly reduced in perivascular, stromal and endothelial cells of EcE. In parallel, perivascular cells exhibited a high expression of estrogen receptor 2 and *HSD17B8* gene that encodes for protein converting estrone (E1) to estradiol (E2), while in endothelial cells *HSD17B2* gene coding for enzyme converting E2 to E1 was downregulated. Overall, our results identified different expression patterns of energy metabolic pathways and steroidogenesis-related genes in perivascular, stromal, and endothelial cells in EcE compared with EuE. Perivascular cells, known to contribute to the restoration of endometrial stroma and angiogenesis, can be a potential target for non-hormonal treatment of endometriosis.

## Introduction

Endometriosis is a chronic, steroid-dependent condition that affects approximately 10% of reproductive-aged women^1^. Endometriosis is characterised by the ectopic growth of endometrial-like tissue, which exerts inflammatory, angiogenic, proliferative and fibrotic processes^1–3^. In 80% of cases endometriotic lesions are localized superficially on the peritoneum^3^. To understand the alterations in cellular status that facilitate cell growth and survival at peritoneal sites, it is essential to closely examine the lesions and the paired endometrial samples from the same women affected by endometriosis at single-cell resolution. Recent single-cell RNA sequencing (scRNA-seq) studies revealed the heterogeneous cellular composition of endometrium, as well as endometriotic lesions, and their microenvironment^4–6^.

Cellular proportions and cell-specific transcriptomic profiles of endometrium change across the menstrual cycle under the control of endocrine factors^7^. Steroid hormones define the phase of the menstrual cycle and lead to morphological and transcriptomic changes in endometrial cells that reflect underlying processes, such as cell proliferation and angiogenesis^8^. These processes require an increase in energy production and substrate biosynthesis^9^. Steroid hormones also activate cell cycle regulators^10–12^ that in turn regulate metabolic reprogramming^13^. The ability of a cell to transit through the cell cycle in response to tissue environmental cues is also dependent on adequate nutrient supply and its metabolic activity and plasticity, referred as adaptive metabolic response^14^.

The endometrium of healthy fertile women is finely controlled by ovarian hormones, and metabolic signaling pathways regulate cellular homeostasis^15^. However, hormonal imbalances, hypoxia and elevated oxidative stress, may affect endometrial tissue processes. A hypoxic microenvironment and altered levels of estradiol (E2) and progesterone have been observed in the ectopic endometrial tissue from women with endometriosis^16–18^, and they can alter cellular metabolism via the regulation of metabolic pathways. Hypoxic microenvironment was also shown to affect metabolic activity in endometriosis^19^. Previous *in vitro* studies reported metabolic switch in endometriosis from oxidative to glycolytic metabolism^19–21^. This shift resembles the Warburg effect, a mechanism actively employed by cancer cells, where cells use aerobic glycolysis to divide and produce biomass and energy at a higher rate^22,23^. Several studies have discussed new strategies of nonhormonal endometriosis management by addressing cellular metabolism^24–26^.

In our study, we investigated different cell types and their metabolic networks in peritoneal lesions (ectopic endometrium, EcE) compared to eutopic endometrium (EuE) using scRNA-seq analysis. This knowledge may facilitate understanding of the metabolic plasticity of EcE in maintaining its homeostasis and growth in a context of unfavorable microenvironment.

## Results

The study was designed as depicted in Fig. 1A. A total of 16,924 cells were obtained from the single-cell transcriptomic analysis of eight paired samples of EcE and EuE from four women with confirmed endometriosis. After quality control and removal of doublets, 7,279 and 7,538 cells from EuE and EcE, respectively, remained for further analysis. The average number of genes per cell was 2,480 (range 1,762 - 3,199).

**Figure 1.**
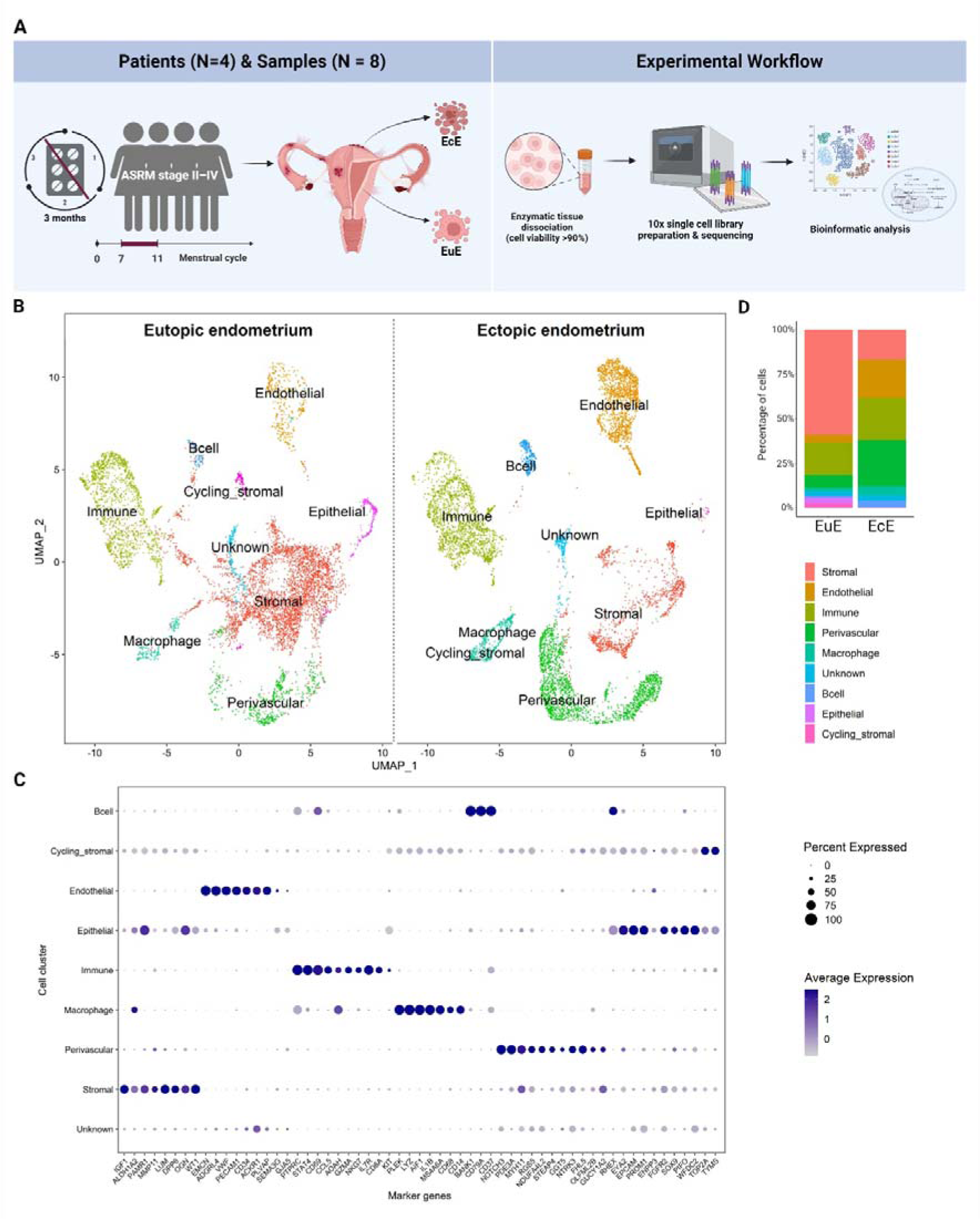
An overview of the single cell landscape of peritoneal endometriosis. **A**. Illustration of the study design. Paired ectopic endometrium (EcE) and eutopic endometrium (EuE) tissue samples from four women with endometriosis during the proliferative phase of menstrual cycle were cryopreserved and thawed, followed by enzymatic tissue dissociation. The isolated cells were submitted for 10x Genomics single cell library preparation and RNA sequencing. Further bioinformatic analyses of cell clusters and metabolic changes were applied. Created with BioRender.com. **B**. Uniform Manifold Approximation and Projection (UMAP) plot of nine major cell clusters. The clusters of EuE – on the left side, EcE – on the right side. **C**. Expression of marker genes in nine major cell clusters, the size of the dots shows the percentage of cells expressing the genes within the clusters, and the colour of the dot corresponds to the average gene expression. Each cluster exhibited the expression of cell-type representative marker genes. **D**. Bar plots representing the relative ratios of 9 major cell populations in EcE (right column) and EuE (left column). Perivascular, endothelial, and immune cell clusters were larger in EcE compared to EuE, while stromal cell cluster was larger in EuE.

### EcE is enriched for perivascular, endothelial, and immune cells

The cells in EuE and EcE were annotated and initially grouped into 16 cell clusters according to the expression of cell type-specific marker genes (Suppl. Fig. 1), followed by re-clustering into nine major cell populations of stromal, endothelial, perivascular, immune, epithelial, cycling stromal cells, macrophages, B cells, and unknown cell population (Fig. 1B). Each cluster was determined based on the differentially expressed genes (DEGs) in that cluster versus overall expression in the rest of the clusters (Suppl. Fig. 2 and Fig. 1C). Cycling stromal cells comprised a population of fibroblasts that expressed cell cycle genes and clustered separately from the main stromal cell population. The unknown cluster included the cells that did not express marker genes that could assign them to a particular cluster.

The statistical analysis of cell proportions between EcE and EuE revealed a significant difference in all nine cell populations, as shown on Fig. 1D and Suppl. Table 1. In EuE, the stromal, epithelial and cycling stromal cell clusters were larger compared to EcE (p < 1.13×10^-103^, p = 1.03×10^-93^, and p = 1.13×10^-103^, respectively), while in EcE, the perivascular, endothelial, immune, macrophage, B cell and unknown cell populations were larger than in EuE (p < 1.13×10^-103^, p < 1.13×10^-103^, p = 5.56×10^-33^, p = 5.93×10^-50^, p = 7,39×10^-63^ and p = 2.62×10^-4^, respectively). Among 9 cell clusters, stromal, perivascular, endothelial and immune clusters were the largest in size in both EuE and EcE (Fig. 1D). The epithelial cell cluster comprised a small cell population in both EuE and EcE, probably due to the cell loss during the tissue dissociation procedure.

### Metabolic reprogramming of perivascular, stromal and endothelial cell populations in EcE

To explore metabolic activity in EcE and EuE, we analyzed the gene expression of 12 metabolic pathways: two regulatory pathways for cellular metabolism, proliferation and angiogenesis (AMP-activated protein kinase (AMPK) signaling, Hypoxia-Inducible Factor-1 (HIF-1) signaling), and 10 catabolic and/or anabolic metabolic pathways (pyruvate metabolism, oxidative phosphorylation (OXPHOS), pentose phosphate pathway, glycolysis/gluconeogenesis, citrate cycle (tricarboxylic acid cycle, TCA), fatty acid (FA) degradation, glutathione metabolism, alanine, aspartate and glutamate (Ala, Asp and Glu) metabolism, FA biosynthesis, and FA elongation).

Among nine cell clusters, perivascular, stromal, and endothelial cell populations had the largest proportions of statistically significant DEGs calculated to the total number of genes in each pathway in a comparison of EcE vs EuE (Suppl. Table 2). Immune, epithelial, cycling stromal and unknown cell clusters had a small number of metabolic DEGs, while macrophages and B cells exhibited no significant difference between EcE and EuE. In perivascular cells, FA elongation (12/27, 44.4%), FA degradation (18/43, 41.9%), pentose phosphate pathway (11/30, 36.7%), HIF-1 signaling (39/109, 35.8%), and pyruvate metabolism (16/47, 34.0%), exhibited the highest percentages of DEGs. In stromal cell population, AMPK signaling (35/121, 28.9%), FA elongation (7/27, 25.9%), pyruvate metabolism (12/47, 25.5%), FA degradation (10/43, 23.3%), and glutathione metabolism (13/57, 22.8%) were among the top altered pathways. While in endothelial cells, glutathione metabolism (12/57, 21.1%), AMPK signaling (23/121, 19%), HIF-1 signaling (21/109, 19.3%), Ala, Asp and Glu metabolism (7/30, 18.9%), and FA biosynthesis (3/18, 16.7%) had the highest proportions of DEGs.

Next, we applied single-cell pathway analysis (SCPA) analysis to evaluate the activity of 12 metabolic pathways in all cell clusters together. According to this analysis, a higher Q value refers to a higher difference in pathways between EcE and EuE, considered as two conditions. The analysis ranked HIF-1 and AMPK signaling pathways at the top of the list with Q values 6.2 and 6.0, respectively (Fig. 2A). We also compared the activity of metabolic pathways in each cell cluster individually. Similarly, HIF-1 and AMPK signaling pathways were top-ranked across all cell clusters (Suppl. Table 3). Perivascular, stromal, and endothelial cells showed the highest Q values for most of the pathways compared to other cell clusters (Fig. 2B), indicating high pathway differences between EcE and EuE.

**Figure 2.**
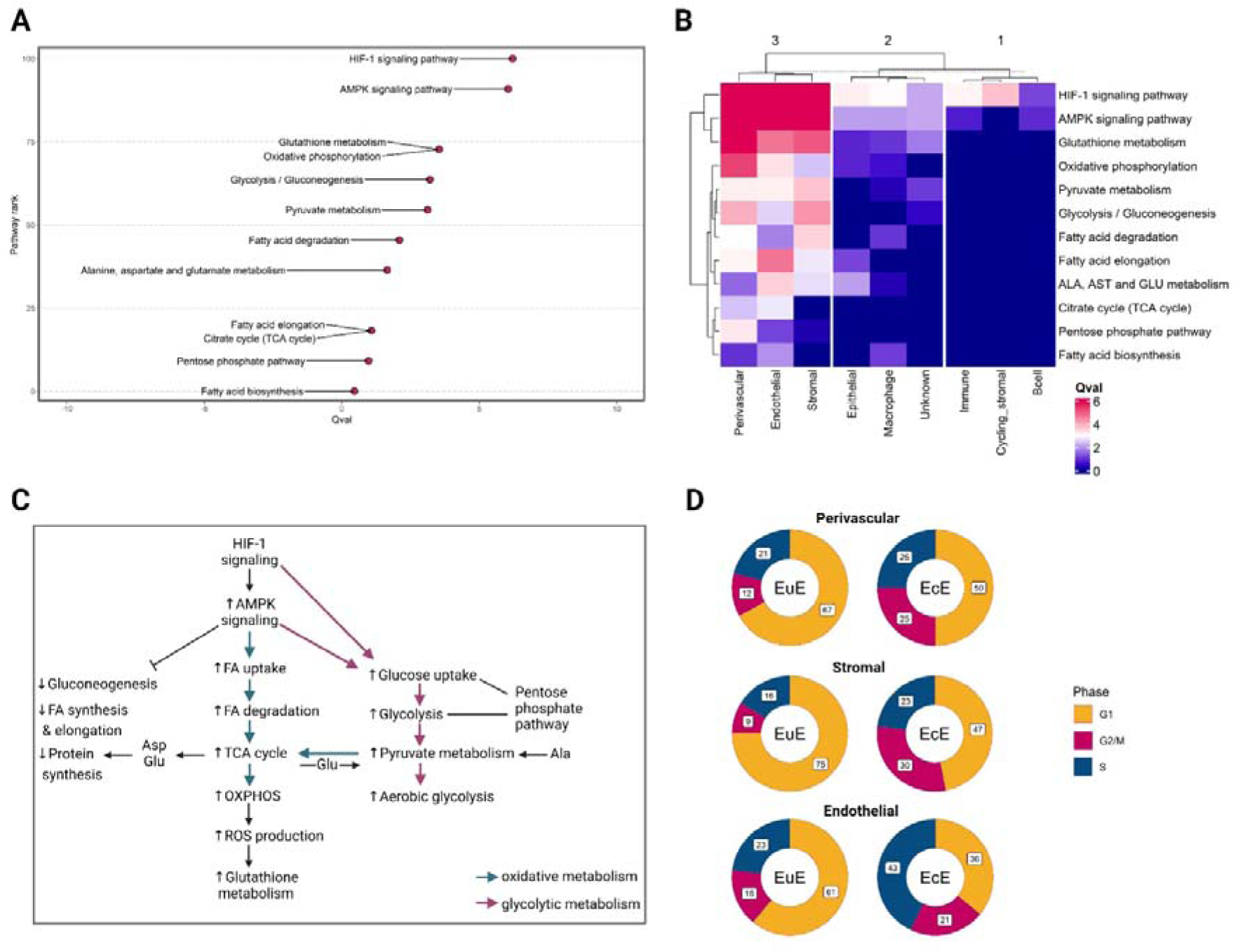
**A**. Ranking of metabolic pathways according to Qvals (Q value) in merged clusters of ectopic endometrium (EcE) compared to eutopic endometrium (EuE). Q value refers to a pathway difference between EcE and EuE, considered as two conditions. A higher number on y axis corresponds to a higher rank of the pathway. **B**. A heatmap showing the Q values of 12 metabolic pathways across 9 cell clusters of EcE compared to EuE. The lower Q value is depicted in blue, the higher Q value in red. **C**. Interconnection between metabolic pathways involved in cellular metabolism. HIF-1 signaling and AMPK signalling are regulatory pathways for the metabolic pathways involved in glycolytic and oxidative metabolism, biosynthesis of macromolecules and nucleic acids. Pink arrows represent glycolytic way of metabolism, blue arrows represent oxidative metabolic pathways. **D**. The proportions of perivascular, stromal and endothelial cells in cell cycle phases in EuE (on the left side) and EcE (on the right side). G1 – cell growth phase (in yellow), S – DNA synthesis phase (in blue), and G2/M - checkpoint and mitosis phase (in red). The numbers in white squares correspond to the percentages of the cells in a given cell cycle phase. Figure 2C was created with BioRender.com.

For further analysis, we focused on three clusters of endothelial, perivascular, and stromal cells that exhibited the highest differences in the regulation of metabolic pathways based on gene expression and SCPA analyses between EcE and EuE. We checked the percentages of statistically significant differentially expressed up- and down-regulated genes in perivascular, stromal and endothelial clusters across 12 metabolic pathways (Table 1). The analysis of differential regulation was performed by comparing EcE to EuE, and the reported statistically significant DEGs are given accordingly (Suppl. Table 4).

**Table 1.** Numbers of up- and down-regulated genes of metabolic pathways in perivascular, endothelial, and stromal clusters of ectopic endometrium compared to eutopic endometrium.

|  |  | Cluster, N (%)<sup>a</sup> of upregulated (UP) or downregulated (DOWN) genes |  |  |  |  |  |
| --- | --- | --- | --- | --- | --- | --- | --- |
|  |  | Perivascular |  | Stromal |  | Endothelial |  |
| Metabolic Pathways | Total N of genes in a pathway | UP, N (%) | DOWN, N (%) | UP, N (%) | DOWN, N (%) | UP, N (%) | DOWN, N (%) |
| Ala, Asp and Glu metabolism | 37 | 5 (13.5%) | 4 (10.8%) | 1 (2.7%) | 1 (2.7%) | 4 (10.8%) | 3 (8.1%) |
| AMPK signaling pathway | 121 | 21 (17.4%) | 18 (14.9%) | 18 (14.9%) | 17 (14.0%) | 11 (9.1%) | 12 (9.9%) |
| Citrate cycle (TCA cycle) | 30 | 9 (30%) | 1 (3.3%) | 3 (10.0%) | 2 (6.7%) | 1 (3.3%) | 2 (6.7%) |
| Fatty acid biosynthesis | 18 | 3 (16.7%) | 1 (5.6%) | 1 (5.6%) | 0 | 2 (11.1%) | 1 (5.6%) |
| Fatty acid degradation | 43 | 15 (34.9%) | 3 (7%) | 10 (23.3%) | 0 | 2 (4.7%) | 0 |
| Fatty acid elongation | 27 | 6 (22.2%) | 6 (22.2%) | 4 (14.8%) | 3 (11.1%) | 2 (7.4%) | 1 (3.7%) |
| Glutathione metabolism | 57 | 7 (12.3%) | 9 (15.8%) | 8 (14%) | 5 (8.8%) | 4 (7%) | 8 (14%) |
| Glycolysis/ Gluconeogenesis | 67 | 19 (28.4%) | 2 (3%) | 10 (15%) | 2 (3%) | 4 (6.0%) | 3 (4.5%) |
| HIF-1 signaling pathway | 109 | 20 (18.3%) | 19 (17.4%) | 13 (11.9%) | 9 (8.3%) | 13 (11.9%) | 8 (7.3%) |
| Oxidative phosphorylation | 134 | 32 (23.9%) | 0 | 5 (3.7%) | 5 (3.7%) | 3 (2.2%) | 2 (1.5%) |
| Pentose phosphate pathway | 30 | 8 (26.7%) | 3 (10.0%) | 4 (13.3%) | 1 (3.3%) | 1 (3.3%) | 2 (6.7%) |
| Pyruvate metabolism | 47 | 16 (34%) | 0 | 12 (25.5%) | 0 | 3 (6.4%) | 2 (4.3%) |

<sup>a</sup>The percentages of the genes are calculated to the total N of genes in a given pathway.

Metabolic pathways are interconnected, and some enzymes with multiple functions play a role in different but related processes (Fig. 2C). As a result, certain genes encoding these enzymes are present in several pathways as discussed below. The DEGs (and their corresponding log_2_fold change, log_2_FC) of metabolic pathways in perivascular, endothelial, and stromal cell clusters of ectopic endometrium compared to eutopic endometrium are presented in Table 2.

**Table 2.**
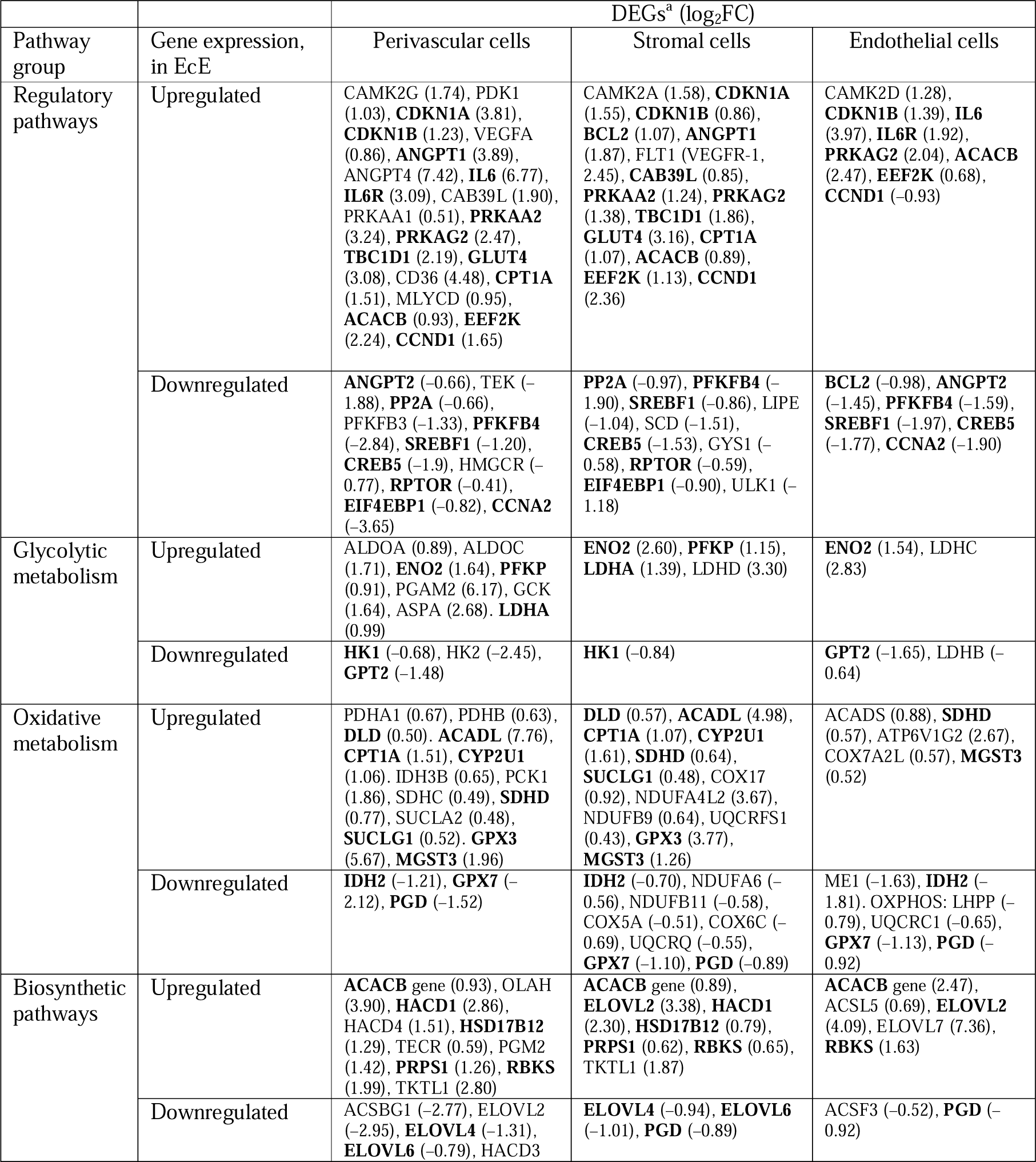
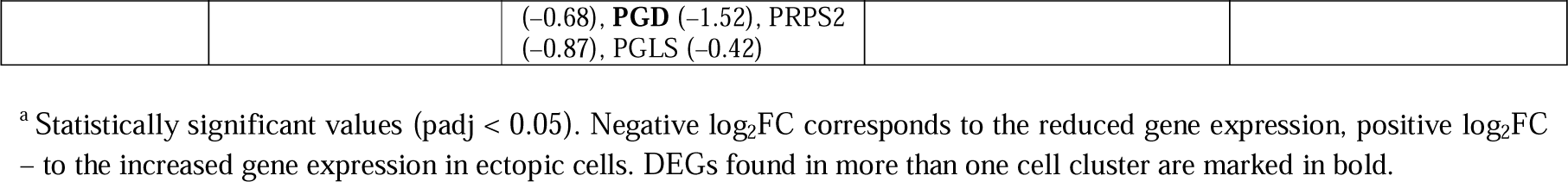
Differentially expressed genes (DEGs) of metabolic pathways in perivascular, endothelial, and stromal cell clusters of ectopic endometrium compared with eutopic endometrium.

### Regulatory metabolic pathways (AMPK signaling pathway and HIF-1 signaling pathway)

AMPK and HIF-1 signaling pathways exhibited a differential regulation of several genes involved in cellular processes like metabolism, proliferation, and angiogenesis. We found a transcriptomic activation of AMPK via HIF-1 target genes, *CAMK2A*, *CAMK2D* and *CAMK2G*, upregulated in all three cell clusters of EcE (Table 2). Apart from that, perivascular and stromal cells in EcE had a high expression of *CAB39L* that activates AMPK. In turns, in perivascular and stromal cells there was a low expression of *PP2A* gene that has an inhibitory activity on AMPK. *PRKAA1*, *PRKAA2* and *PRKAG2* genes encoding subunits of AMP protein kinase were overexpressed in all three clusters of EcE.

AMPK and HIF-1 signaling pathways regulate oxidative and glycolytic metabolism in cells. AMPK signaling activates energy-producing pathways that utilize glucose and FAs as main substrates. In perivascular and stromal cell clusters, we found an upregulation of *TBC1D1* and *GLUT4* genes leading to increased glucose uptake. *PFKFB3* (in perivascular cells) and *PFKFB4* (in perivascular, stromal, and endothelial cells) genes, associated with AMPK-mediated activation of glycolysis and glucose flux in pentose phosphate pathway, were downregulated in EcE. However, AMPK target genes, *CD36* (in perivascular cells), *CPT1A* (in perivascular and stromal cells), and *MLYCD* (in perivascular cells), involved in FA uptake, transport to mitochondria and β-oxidation were upregulated. In addition, in all three clusters of EcE, there was an overexpression of *ACACB* gene converting acetyl-CoA to malonyl-CoA that can be further used for FA synthesis. At the same time, in ectopic perivascular cells, we identified an upregulation of *PDK1*, a HIF-1 target gene that inhibits oxidative metabolism. Whereas, HIF-1 target genes encoding key glycolytic enzymes like *ALDOA*, *ENO2*, *LDHA* were overexpressed. As these genes are common for HIF-1 signaling as well as for glycolysis, pyruvate metabolism and TCA cycle, we describe the differential expression of corresponding genes within those pathways.

Another AMPK-mediated effect is an inhibition of biosynthetic processes. In our analysis, we found that the genes associated with biosynthetic metabolism were both up- and downregulated in EcE. For example, the above-mentioned *ACACB* gene was upregulated, while *SREBF1* (in perivascular, stromal, and endothelial cells), *LIPE* (in stromal cells) and *SCD* (in stromal cells) that are associated with FA and sterol synthesis, were downregulated. Other genes that are key regulators and activators of gluconeogenesis (*CREB5* in perivascular, stromal, and endothelial cells), glycogen synthesis (*GYS1* in stromal cells), and sterol synthesis (*HMGCR* in perivascular cells) were inhibited in EcE. There was a downregulation of *RPTOR* (in perivascular and stromal cells) gene encoding a protein from mTORC1 complex and involved in the activation of protein synthesis via phosphorylated EIF4EBP1. *EEF2K*, an AMPK-mediated inhibitor of protein synthesis, was overexpressed in all three clusters of EcE.

AMPK and HIF-1 signaling pathways are known to regulate cell cycle, proliferation, and survival. *CDKN1A* (in perivascular and stromal cells) that has an inhibitory effect on cyclin-dependent kinases, and *CDKN1B* (in perivascular, stromal cell and endothelial cells) involved in cell cycle arrest at G1 phase were upregulated in EcE. Additionally, we observed an inhibition of *CCNA2* gene (in perivascular and endothelial cells of EcE) encoding cyclin A2, which is known to be overexpressed in S phase of cell cycle and involved in cell cycle transition. However, *CCND1* gene that plays a critical role in G1-S transition was upregulated in perivascular and stromal cells but downregulated in endothelial cells. In addition, in EcE we found an upregulation of anti-apoptotic gene *BCL2* in stromal cells and its downregulation in endothelial cells. In stromal cells of EcE, we also observed a downregulation of *ULK1* gene that mediates autophagy.

We found a differential expression of HIF-1-mediated genes involved in angiogenesis in EcE compared to EuE. *VEGFA* (in perivascular cells), *ANGPT1* (in perivascular and stromal cells) gene that plays a role in vessel growth, and *ANGPT4* (in perivascular cells) important for interactions between perivascular and vascular endothelial cells, were upregulated, while *ANGPT2* (in perivascular and endothelial cells), known as an antagonist of *ANGPT1*, and *TEK* receptor (in perivascular cells), were downregulated. Additionally, there was an overexpression of *IL6* (in perivascular and endothelial cells) that plays a role in the activation of *VEGF*, its receptor *IL6R* (in perivascular and endothelial cells), and VEGFR-1 encoding gene *FLT1* (in stromal cells).

### Glycolytic metabolism

Glycolysis pathway is responsible for pyruvate production and is regulated by both HIF-1 and AMPK signaling pathways. In EcE, we found an overexpression of key glycolysis-related genes like *ALDOA* and *ALDOC* (in perivascular cells), *ENO2* (in perivascular, stromal and endothelial cells), *PFKP* (in perivascular and stromal cells), and *PGAM2* (in perivascular cells). Interestingly, we observed a downregulation of *HK1* (in perivascular and stromal cells), and *HK2* (in perivascular cells) that regulate the first step of glycolysis, however in perivascular cells there was an upregulation of *GCK* that takes the role of hexokinases to phosphorylate glucose and thereby maintains glycolysis.

Ala, Asp and Glu glucogenic amino acids can be used by the cells to produce glycolysis intermediates in the energy depleted state of the cells. Additionally, there is a reverse reaction, when Ala can be synthesized from the intermediates of glycolysis (pyruvate), and Asp and Glu can be converted from TCA intermediates. In perivascular and endothelial cells of EcE, we found a differential regulation of genes related to glucogenic amino acid metabolism with some genes upregulated, for example *ASPA* (in perivascular cells) involved in Asp and acetate production, and other genes like *GPT2* (in perivascular and endothelial cells), which catalyzes pyruvate production from alanine, being downregulated. Overall, the gene expression of this pathway in stromal cells was similar between EcE and EuE.

In EcE, we found a differential expression of lactate dehydrogenases (LDH) that convert pyruvate to L- and D-lactate. In perivascular cells, differentially expressed *LDH* genes were represented by *LDHA*, in stromal cells by *LDHA* and *LDHD*, and in endothelial cells by *LDHB* and *LDHC*. This refers to a catabolic way of pyruvate utilization in cells, called aerobic glycolysis via LDH enzyme regulation. However, apart from the genes related to the pyruvate conversion to lactate, in EcE we found an upregulation of enzymes involved in the conversion of pyruvate to acetyl-CoA. *PDHA1*, *PDHB* and *DLD* genes, which encode enzymes from pyruvate dehydrogenase complex that links glycolysis and TCA cycle, were upregulated in perivascular and stromal cells. Pyruvate can be produced from other substrates apart from glucose, and in ectopic endothelial cells we found a downregulation of *ME1* gene encoding an enzyme that converts malate (TCA cycle intermediate) to pyruvate.

### Oxidative metabolism

With the analysis of genes from FA degradation pathway, in EcE we identified an upregulation of genes related to the formation of acetyl-CoA from FAs, called β-oxidation. FAs are one of the main sources of energy, and acetyl-CoA within the cells is further used in oxidative metabolism. Several enzymes that catalyze the steps of FA β-oxidation in mitochondria were upregulated, for example *ACADL*, *CPT1A*, and *CYP2U1* in perivascular and stromal cells, and *ACADS* in endothelial cells.

In EcE, we found an upregulation of genes encoding key proteins of TCA cycle in mitochondria, where oxaloacetate and acetyl-CoA produced from FAs and/or pyruvate are used as a main substrate. Among these genes, *IDH3B*, *PCK1*, *SDHC* and *SUCLA2* were overexpressed in perivascular cells, *SUCLG1* in perivascular and stromal clusters, and *SDHD* in all three clusters. *IDH2*, involved in TCA cycle with glutamine as a substrate, was downregulated in all three clusters, probably indicating a higher use of FAs and pyruvate over glutamine in TCA cycle.

NADH and FADH (the products of TCA cycle, FA oxidation and glycolysis) are further utilized in electron transport chain consisting of OXPHOS enzyme complexes where ATPs are produced. In ectopic perivascular cells, we identified an overexpression of 24% of genes, encoding five multiprotein complexes of the electron transport chain of OXPHOS. In stromal and endothelial cells, OXPHOS pathway was not remarkably altered as in perivascular cluster, probably being similarly regulated and utilized in both EcE and EuE. Stromal cells exhibited a downregulation of *NDUFA6*, *NDUFB11*, *COX5A*, *COX6C*, and *UQCRQ*, and upregulation of *COX17*, *NDUFA4L2*, *NDUFB9*, *SDHD*, and *UQCRFS1* that encode for OXPHOS complexes. In endothelial cells, *ATP6V1G2*, *COX7A2L*, and *SDHD*, were upregulated, and *LHPP* and *UQCRC1* were less expressed in EcE compared to EuE.

We found a differential regulation of glutathione peroxidases, reductases, transferases and other genes that are involved in glutathione synthesis and neutralization of oxygen reactive species that are mainly formed during oxidative metabolism. Some genes like *GPX3* (in perivascular and stromal cells, respectively) and *MGST3* (in perivascular, stromal cell and endothelial cells) were higher expressed in EcE, while other genes like *GPX7* (in ectopic perivascular, stromal cell and endothelial cells) and *PGD* (in ectopic perivascular, stromal cell and endothelial cells) were overexpressed in EuE.

### Biosynthetic metabolism

Regarding FA synthesis pathway, we identified an overexpression of *ACACB* gene (in perivascular, stromal, and endothelial cells) that produces malonyl-CoA, a driver of FA synthesis, from acetyl-CoA (mentioned in AMPK signaling) in EcE. However, the *ACSF3* gene that also synthesizes malonyl-CoA from malonic acid was not differentially regulated in perivascular and stromal cells between EcE and EuE, and in endothelial cells it was even less expressed in EcE compared to EuE. *OLAH* gene that participates in FA release from FA synthase was upregulated in perivascular cells of EcE. The genes involved in the conversion of FA to acyl-CoA, which is utilized in FA elongation process, were downregulated in perivascular cells (*ACSBG1*) and upregulated in endothelial cells (*ACSL5*) of EcE.

FA elongation pathway, consisting of four steps for long-chain FA synthesis from malonyl-CoA and acyl-CoA, was also differentially regulated in EcE. The genes encoding enzymes that catalyze the rate-limiting first and third steps, were up- and downregulated in perivascular cells (*ELOVL2*, *ELOVL4*, *ELOVL6*, *HACD1*, *HACD3*, *HACD4*), and in stromal cells (*ELOVL2*, *ELOVL4*, *ELOVL6*, *HACD1*), and upregulated in endothelial cells (*ELOVL2* and *ELOVL7*). While the genes from the second and forth steps were upregulated in perivascular (*HSD17B12* and *TECR*), and stromal cells (*HSD17B12*).

Pentose phosphate pathway and glycolysis are linked pathways and depending on energy need, a cell can employ one or another pathway. If a cell does not require additional energy synthesis, the glucose enters pentose phosphate pathway to produce nucleotides and NADPH that can further be used in anabolic processes. In EcE, we observed an upregulation of genes involved in ribose and nucleotide synthesis and metabolism, like *PGM2* in perivascular cells, *PRPS1* and *TKTL1* in perivascular and stromal cells, and *RBKS* in all three clusters. However, *PGD* (in perivascular, stromal and endothelial cells), *PRPS2* and *PGLS* (in perivascular cells) genes, involved in oxidative steps of the pathway, were downregulated in EcE.

### Progesterone resistance and elevated estradiol signaling genes in EcE

As cellular metabolism in endometrium is partially regulated by steroid hormones, we next examined the expression levels of steroidogenic enzymes between EcE and EuE. The differential expression analysis revealed a significantly lower expression of *PGR* in perivascular and endothelial cells of EcE (Table 3). The *ESR2* and *HSD17B8* genes involved in the activation of estrogen target genes and conversion of a less potent estrogen E1 to E2, respectively, were significantly upregulated in ectopic perivascular cells. The expression level of *ESR1* was significantly decreased in EcE compared to EuE. A significant downregulation of *SRD5A3, SRD5A1* and *AR* genes related to androgen receptor and androgen production, as well as *CYP11A1* and *HSD11B2* genes related to the synthesis and metabolism of steroids and cortisol, respectively, was observed in perivascular cell population in EcE. Stromal cells of EcE overexpressed *AKR1C1* and *AKR1C2* genes involved in the metabolism of progesterone, *HSD11B1* gene that encodes an enzyme producing cortisol, and *HSD17B6* and *HSD17B11* that enhance androgen metabolism. Endothelial cells also exhibited the upregulation of genes related to the progesterone (*AKR1C1*) and ketones (*HSD17B11*) metabolism. Together, the results of steroidogenesis analysis showed transcriptomic alterations of progesterone signalling and increased resistance (*PGR, AKR1C1, AKR1C2*), and elevated estrogen activity by the increased expression of genes involved in synthesis of E2, reduced E2 conversion/inactivation into E1, downregulation of ERα and the overexpression of ERβ (*HSD17B8, HSD17B2* and *ESR2*, respectively). According to previous reports on E2 role in endometriosis (Rižner, 2009), this might refer to the increased proliferative activity of cells in endometriotic lesions. To investigate the portion of proliferating cells in perivascular, stromal and endothelial cells of EcE, we next checked proportions of cell clusters in cell cycle phases in EcE and EuE.

**Table 3.**
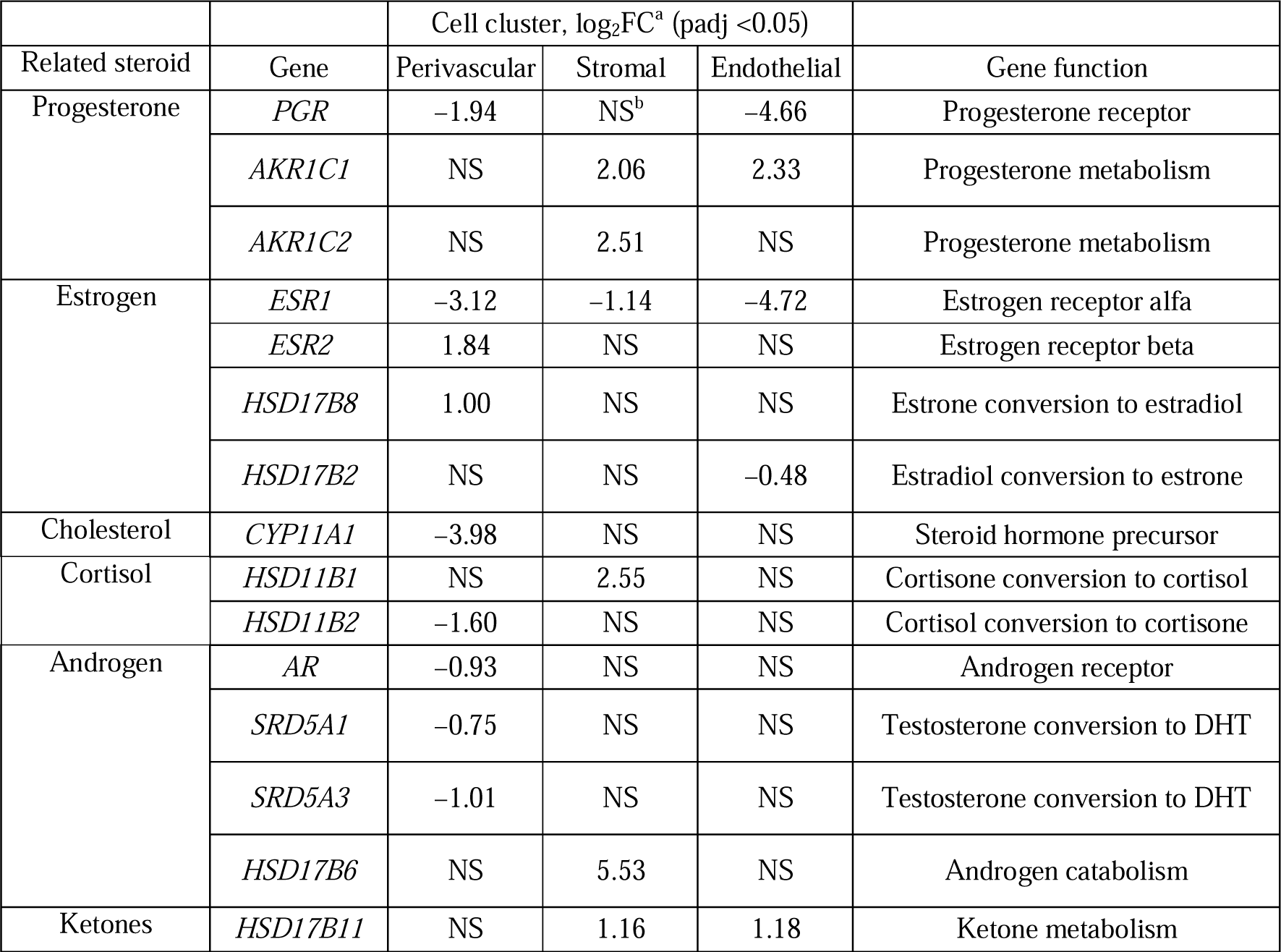
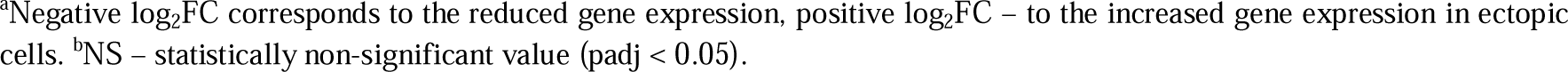
Statistically significant differentially expressed genes coding for enzymes involved in the regulation of synthesis, conversion and metabolism of steroids in perivascular, endothelial, and stromal cell clusters of ectopic endometrium compared with eutopic endometrium.

|  |  | Cell cluster, log<sub>2</sub>FC<sup>a</sup> (padj <0.05) |  |  |  |
| --- | --- | --- | --- | --- | --- |
| Related steroid | Gene | Perivascular | Stromal | Endothelial | Gene function |
| Progesterone | *PGR* | -1.94 | NS<sup>b</sup> | -4.66 | Progesterone receptor |
|  | *AKR1C1* | NS | 2.06 | 2.33 | Progesterone metabolism |
|  | *AKR1C2* | NS | 2.51 | NS | Progesterone metabolism |
| Estrogen | *ESR1* | -3.12 | -1.14 | -4.72 | Estrogen receptor alfa |
|  | *ESR2* | 1.84 | NS | NS | Estrogen receptor beta |
|  | *HSD17B8* | 1.00 | NS | NS | Estrone conversion to estradiol |
|  | *HSD17B2* | NS | NS | -0.48 | Estradiol conversion to estrone |
| Cholesterol | *CYP11A1* | -3.98 | NS | NS | Steroid hormone precursor |
| Cortisol | *HSD11B1* | NS | 2.55 | NS | Cortisone conversion to cortisol |
|  | *HSD11B2* | -1.60 | NS | NS | Cortisol conversion to cortisone |
| Androgen | *AR* | -0.93 | NS | NS | Androgen receptor |
|  | *SRD5A1* | -0.75 | NS | NS | Testosterone conversion to DHT |
|  | *SRD5A3* | -1.01 | NS | NS | Testosterone conversion to DHT |
|  | *HSD17B6* | NS | 5.53 | NS | Androgen catabolism |
| Ketones | *HSD17B11* | NS | 1.16 | 1.18 | Ketone metabolism |

<sup>a</sup>Negative log<sub>2</sub>FC corresponds to the reduced gene expression, positive log<sub>2</sub>FC – to the increased gene expression in ectopic cells. <sup>b</sup>NS – statistically non-significant value (padj < 0.05).

### Higher proportions of dividing cells in EcE

As we observed changes in metabolic activity and steroidogenesis in perivascular, stromal and endothelial cell clusters of EcE, we examined proportions of these three cell types by cell cycle phases in EcE and EuE. The analysis was based on the expression of cell cycle genes corresponding to three cell cycle phases: G1 phase of cell growth, S phase of DNA synthesis, and G2/M phase (combined G2 and M phases) of checkpoint and cell division (Suppl. Table 5).

In perivascular, stromal, and endothelial cell populations of EuE over 60% of the cells were in the G1 phase, 16% to 23% accounted for the S phase, and only 9% to 16% of the cells were in the G2/M phase of the cell cycle (Fig. 2D).

In contrast, in EcE, less cells were detected in the G1 phase (36% to 50%), and higher proportions of cells were in G2/M (21% to 30%) and S (23% to 43%) phases compared to EuE. The differences between ectopic and eutopic cell proportions were significant for each cluster in the three phases according to Fisher’s exact test (Suppl. Table 5), except for stromal cells in G2/M phase. These data suggest a higher proliferation rate for EcE cells, probably associated with altered steroidogenesis as described in the previous subsection.

### Differential expression and pathway analyses revealed altered proliferation, apoptosis, migration and angiogenesis processes in EcE

Additionally, we checked the overall differential gene expression profile and associated pathways in perivascular, stromal and endothelial cell populations. The differential expression analysis of genes between EcE and EuE identified 5,602, 3,733 and 3,233 altered genes in perivascular, stromal, and endothelial cell clusters, respectively. We performed Kyoto Encyclopedia of Genes and Genomes (KEGG) pathway analysis of these genes and found the enrichment of extracellular matrix (ECM)–receptor interaction, focal adhesion, PI3K-Akt signaling pathway, cell cycle, proteoglycans in cancer, cell adhesion molecules, AMPK and Hippo signaling pathways, and ribosome pathway altered between EcE and EuE tissues (Suppl. Fig. 3, 4). These pathways are involved in cell–ECM interactions, cellular metabolism, proliferation, cell survival, migration and angiogenesis. Among the top 20 DEGs (Suppl. Fig. 5) we identified genes participating in these processes and previously reported to play a role in endometriosis and other conditions characterised by an increased proliferation and invasive growth. In the perivascular cell compartment of EcE, we identified a significant downregulation of tumor suppressor *PAMR1,* metalloproteinase *ADAM12*, lncRNA *MEG8*, *DIO2* and *VCAN* genes, and an upregulation of antiapoptotic gene *NTRK2*. *SKAP2* was overexpressed in both perivascular and endothelial cell clusters of EcE. *ADIRF*, tumor-associated genes *CLU*, *FAM13C,* as well as *NGF*, *PRELP, RERG, PTGIS, GPC3* genes were among the top 20 overexpressed genes in EcE stromal cell cluster. *AQP1* was found to be upregulated in the endothelial and stromal cells of EcE compared with EuE. Overall, DEG and KEGG analyses showed enrichment of pathways related to cell proliferation, survival, and angiogenesis in EcE.

## Discussion

In this single-cell transcriptomic study, we explored the gene expression from the perspective of cellular metabolism, steroidogenesis and cell cycle, in paired samples of EuE and EcE from women with endometriosis. To sustain cell survival, growth, and angiogenesis, endometriotic cells must be adaptable to adjust their metabolism based on cellular needs and environmental conditions. We found that three cell types of perivascular, stromal, and endothelial cells in EcE had different expression patterns of metabolic pathways compared to EuE (Fig. 2B and Table 1). We also observed a transcriptomic downregulation in progesterone signaling in these three clusters of EcE and an increased expression of genes associated with higher estrogen activity in perivascular cell population of EcE compared with EuE (Table 3). Moreover, in EcE, the proportions of perivascular, stromal and endothelial cells in G2/M and S phases were significantly larger than in EuE (Fig. 2D), indicating active cellular proliferation in endometriotic lesions.

The network of perivascular, stromal and endothelial cell play a significant role in endometrial tissue growth. Stromal cells comprise the most abundant cell type in endometrium that consists of multiple subpopulations^5,30^ and actively interact with neighboring cells. Previous *ex vivo* and *in vitro* endometriosis studies showed the Warburg effect at RNA and protein levels in stromal cells^20,21^. Perivascular cells are represented by two cell types, vascular smooth muscle cells and pericytes that reside in the area around large and small vessels, respectively, and interact with endothelial cells, promoting angiogenesis^29^. Pericytes were shown to act as mesenchymal progenitors and affect regenerative processes of endometrial stroma^30–32^. We found larger populations of perivascular, endothelial, and immune cells of EcE compared with EuE, whereas EuE was enriched with stromal cells (Fig. 1D), similar to the results reported by Zhu et al.^28^. Tan et al (2022). found endothelial cell enrichment in ectopic endometrium and reported a new subpopulation of perivascular cells characteristic to endometriosis, expressing gene markers *MYH11* and *STEAP4*, which we also identified in both EcE and EuE (Suppl. Table 6).

Metabolic pathways are regulated by AMPK and HIF-1 signaling^22^. AMPK pathway is activated in response to energy stress by sensing increases in AMP:ATP and ADP:ATP ratios, leading to the activation of glycolysis and FA catabolism, an inhibition of macromolecular synthesis and cell cycle arrest^33,34^. The AMPK pathway, downregulated by steroid hormones, was shown to be involved in inflammation, metabolic regulation, and apoptotic processes in endometriosis^35^. HIF-1 signaling, a major regulator of oxygen homeostasis in cells, has also an impact on steroidogenesis and cell metabolism to sustain cell growth, angiogenesis and survival with a downstream activation of metabolic pathways in endometriotic lesions^18^. The first step of glycolytic way of energy production is glycolysis, during which glucose is converted into pyruvate with the production of 2 molecules of ATP. In oxidative metabolism, pyruvate is then converted to acetyl-CoA through the link reaction, which then enters the TCA cycle in the mitochondria, followed by OXPHOS. FAs undergoing degradation are another source of acetyl-CoA. During this metabolic process, a cell can produce around 30 molecules of ATP^36^.

Some cells, in the presence of oxygen, switch from oxidative to glycolytic ATP synthesis, and pyruvate is converted into L-lactate or D-lactate by LDH enzymes, with regenerated NAD+ that is further re-used in glycolysis, thereby sustaining this reaction^36^. This metabolic switch is known as the Warburg effect, also referred as aerobic glycolysis, observed in cancer cells and in highly proliferative cells that aims to synthesize biomass and produce energy to maintain cell proliferation^23^. An altered energy metabolism toward aerobic glycolysis with inhibited oxidative respiration was observed in endometriosis studies on primates, and *in vitro* studies of endometriotic lesions, endometriotic stromal cells and mesothelial cells from women with endometriosis^19,20,37,38^. A simultaneous activation of aerobic glycolysis and OXPHOS has been recently identified in cancer cells based on transcriptome data analysis^39,40^, and AMPK and HIF-1 signaling played a key regulatory role in sustaining sufficient energy and macromolecular synthesis in the cells^39^. The role of L-lactate and D-lactate in glycolytic and oxidative metabolism was recently examined^41^. Remarkably, both forms of lactate were demonstrated to enter mitochondria without being metabolized and stimulate TCA cycle and OXPHOS.

In our study, we observed a HIF-1- and AMPK-mediated activation of glycolytic and oxidative metabolic systems in perivascular cells of EcE. We found an upregulation of genes that increase glucose uptake (*TBC1D1* and *GLUT4*), key regulators of glycolysis (*GCK, ENO2, PFKP, PGAM2, ALDOA* and *ALDOC*), and overexpressed *LDHA* that mediates aerobic glycolysis. In parallel, there was an increased expression of PDH complex enzymes (*PDHA1, PDHB* and *DLD*) that convert pyruvate to acetyl-CoA. Additionally, in perivascular cells of EcE we identified an upregulation of genes related to an increased FA uptake and its transport to mitochondria (*CD36, CPT1A* and *MLYCD*), and activation of beta-oxidation of FAs with the production of acetyl-CoA (e.g. *ACADL, CPT1A, CYP2U1*). Moreover, there was an upregulation of the key genes of TCA cycle (*IDH3B, PCK1, SDHC, SDHD, SUCLA2* and *SUCLG1*), and overexpression of 24% of genes encoding five OXPHOS multiprotein complexes. Collectively, these data might refer to the above-mentioned co-activation of aerobic glycolysis and oxidative metabolism in perivascular cells of EcE.

Stromal cells of EcE exhibited overall less genes differentially regulating glycolytic and oxidative metabolism than perivascular cells. We found an upregulation of genes related to glucose uptake (*TBC1D1* and *GLUT4*), key enzymes of glycolysis (*ENO2* and *PFKP*), and downregulation of *HK1.* Similarly to perivascular cells, we observed an overexpression of LDH genes referring to aerobic glycolysis (*LDHA* and *LDHD*), and an upregulation of a PDHA complex enzyme (*DLD*) that associates glycolysis and TCA cycle. We also observed an increased expression of genes regulating FA transport to mitochondria and beta-oxidation (*CPT1A, ACADL, ACADS, CYP2U1*, etc.). In addition, there was an upregulation of two TCA key genes (*SDHD* and *SUCLG1*), and both up- and down-regulation of genes encoding OXPHOS complexes. Interestingly, four of five downregulated genes of OXPHOS complexes (*NDUFA6, COX5A, COX6C* and *UQCRQ*), were also found to be lower expressed at protein level in cultured stromal cells from EcE compared with EuE from women with endometriosis and controls^20^. Among the five upregulated genes that we found in OXPHOS pathway, two of them were downregulated at protein level in cultured stromal cells from EcE^20^. Overall, there was an activation of genes related to both glycolytic and oxidative ways of energy production in stromal cells of EcE, however to a lesser extent when compared with perivascular cells. And we did not observe the distinct transcriptomic signature of Warburg effect in the stromal cells of EcE as previously reported. This discrepancy can be explained by the differences between *ex vivo* and *in vitro* experiments.

In the endothelial cell population of EcE, the gene expression pattern of metabolic pathways was similar to that in perivascular and stromal cells, but less altered. Probably, it can be explained by an analogous process of angiogenesis occurring in both EcE and EuE in the proliferative phase of menstrual cycle. Additionally, we found an AMPK-mediated suppression of biosynthetic processes via key regulators of gluconeogenesis (*CREB5*), FA and sterol synthesis (*ACACB, SREBF1, LIPE, SCD, HMGCR*), protein synthesis (*RPTOR, EIF4EBP1, EEF2K*), and glycogen synthesis (*GYS1*) in perivascular, stromal and endothelial cells of EcE. These results might suggest a more catabolic state of perivascular and stromal cells but not endothelial cells in EcE with co-activation of glycolytic and oxidative metabolism to sustain energy requirements.

Non-hormonal therapy of endometriosis has a potential to disrupt metabolic activity in the cells of EcE^25^. For example, McKinnon et al. (2014) showed a high protein expression of glucose transporter GLUT4 in epithelial and stromal cells of EcE suggesting GLUT4 role in glucose supply for lesions and proposing it as a target for therapy. Other targets for endometriosis management could be PDH and PDK1 that play key roles for oxidative and glycolytic metabolism. Their regulator dichloroacetate was studied by Horne et al. (2019) on mice model of endometriosis demonstrating reduction of endometriotic lesion size and lactate level in peritoneal microenvironment.

Steroid hormones play an important role in the regulation of adaptive metabolic response. The reduction of PGR (progesterone resistance), and an increase of E2 production as well as the prevention of E2 inactivation have been observed in endometriosis^42–45^. E2 regulates the transcription of genes involved in cell proliferation, survival and angiogenesis in EcE and other tissues, for example by activating PI3K/AKT and HIF-1 pathways^46–48^. E2 in endometrium is primarily bound to ERα (coded by *ESR1*), whereas in endometriosis ERβ (coded by *ESR2*) mediates E2 effects and is overexpressed in both EuE and EcE, while ERα is downregulated in EcE^45,49,50^. ERβ was reported to be expressed in mitochondria and play a role in metabolic processes as well as prevention of apoptosis by regulating autophagy-related genes in endometriosis^7,51^. We examined the expression of genes involved in steroidogenesis in stromal, perivascular and endothelial cells and observed a significantly reduced expression of *ESR1* in EcE. *PGR* was significantly downregulated in perivascular and endothelial cells, and an increased expression of genes associated with progesterone metabolism (*AKR1C1* and *AKR1C2*) in stromal and endothelial cells. Perivascular cells exhibited a high expression of *ESR2* and *HSD17B8* gene that codes for protein converting E1 to E2, while in endothelial cells, *HSD17B2* coding for enzyme converting E2 to E1 was downregulated.

Cellular metabolism is a process tightly coupled with the cell cycle due to high demand for energy and substrates in a dividing cell. The stromal cells of EuE at the proliferative phase of menstrual cycle were found to be mostly in the G1 cell cycle phase of cell growth, characterised by active macromolecular synthesis^7,51^. McKinnon et al.^52^ studied isolated endometrial stromal cells from women with and without endometriosis, and found that a large portion of the cells was in G1 phase (70%) and only 15% of the cells were in G2M or S phases, which correlates with our observations in the EuE. We identified 50% or less of perivascular, stromal and endothelial cells of EcE in the G1 phase, and larger cell proportions in S and G2/M phases when compared to EuE. This might refer to a higher proliferative rate in the cells of EcE compared to EuE. Taken together, these data suggest an altered regulation of steroid hormones and downstream target genes of PGR, ERα and ERβ, and the activation of energy metabolic pathways in EcE (Fig. 3).

**Figure 3.**
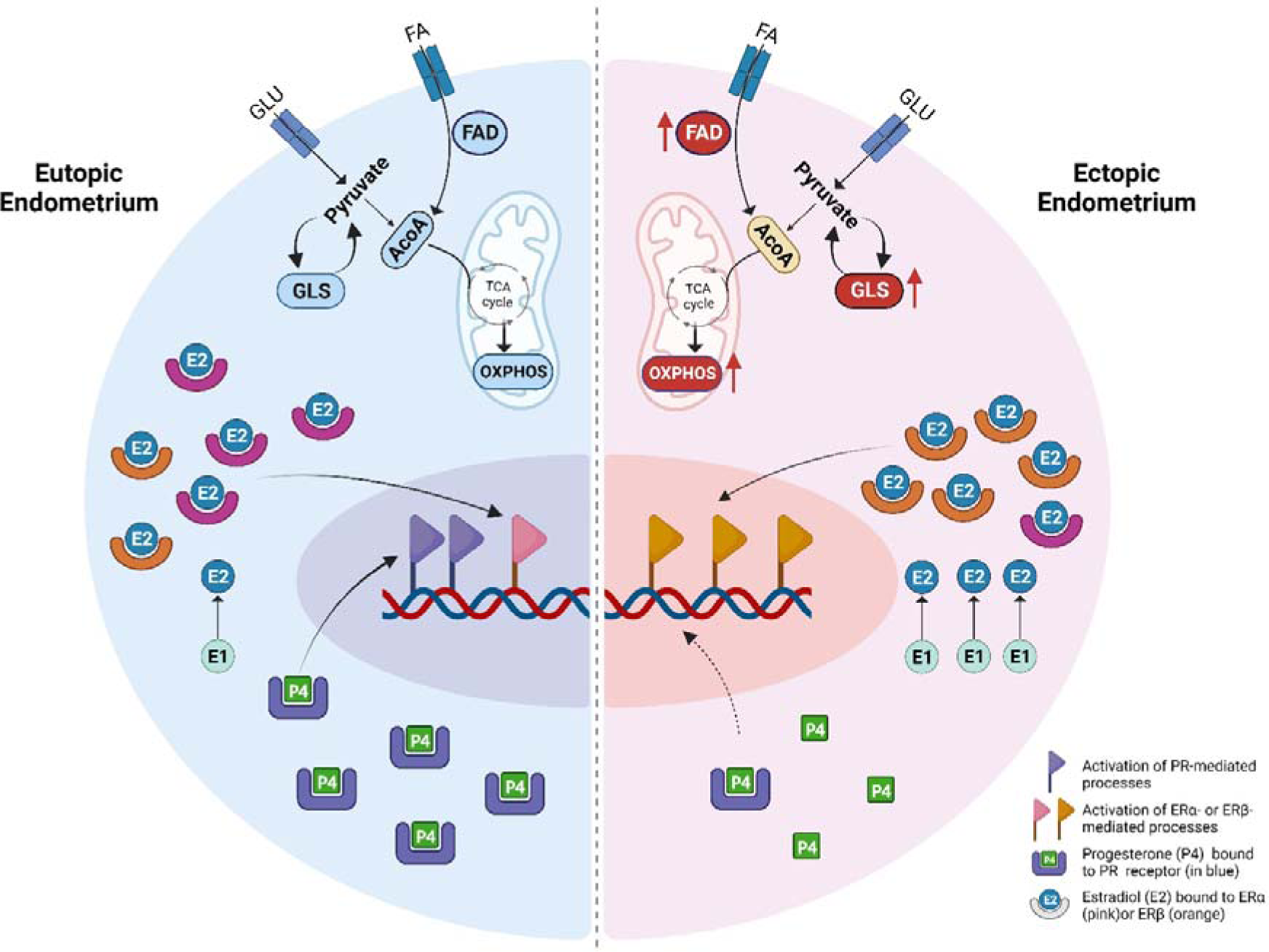
A proposed model of possible interconnected mechanisms of steroid hormone regulation and altered cellular metabolism in perivascular cells of ectopic endometrium (EcE, a half of a cell depicted on the right side) compared to eutopic endometrium (EuE, on the left side). In the EcE the following changes are observed: reduced production of progesterone receptor (PR, in blue) contributing to progesterone (P4) resistance; decreased level of ERα (in pink), increased conversion of estrone (E1) into estradiol (E2), and increased synthesis of ERβ (in orange), leading to a nuclear activation of estrogen target genes; upregulation of genes related to increased glucose and fatty acid (FA) uptake, activated FA degradation (FAD), OXPHOS and glycolysis (GLS). FA – fatty acids, GLU – glucose, ACoA – acetyl-CoA. Created with BioRender.com.

Our scRNA study has some limitations. Firstly, the sample size was limited to four women with endometriosis from the proliferative phase of menstrual cycle; nevertheless, we identified the major cell clusters in both EuE and EcE, and revealed their steroidogenic and metabolic cell type heterogeneity. Moreover, both the samples of EuE and EcE were derived from the same women. These paired samples make the results more robust. Secondly, the study focused exclusively on transcriptomic data analysis with no functional validation of the results. However, previous studies on human cancer have reported that transcriptomic changes in metabolism might refer to and predict metabolic activity in cells^53,54^. Further studies may include a larger and more diverse patients’ population for ethnicity and endometriosis stage, to explore interpatient and interpopulation variability of metabolic reprogramming in endometriotic lesions with functional analysis. Thirdly, we observed a small population of epithelial cells in both EuE and EcE, most likely due to the cell loss during the tissue dissociation procedure. Some previous studies reported smaller clusters of epithelial cells in EcE compared to EuE of women with endometriosis and controls^6,28^.

## Conclusions

Within this study we gained an insight into metabolic heterogeneity, steroidogenesis and cellular proportions in cell cycle phases in EcE and EuE. We found changes in the regulation of progesterone and estrogen signaling pathways in perivascular, stromal and endothelial cells of EcE compared with EuE, which might have a direct effect on cellular metabolism and contribute to cellular proliferation and angiogenesis. The metabolic pathways were differentially regulated in perivascular, stromal and to a less extent in endothelial cell clusters, and metabolic pathway activity alterations between EcE and EuE were the highest for AMPK, HIF-1, glutathione metabolism, OXPHOS, and glycolysis/gluconeogenesis.

Remarkably, we identified a transcriptomic co-activation of glycolysis and OXPHOS in perivascular and stromal cells of EcE compared to EuE, possibly to meet the increased energy demands of proliferating cells. Perivascular cells, known to contribute to the restoration of endometrial stroma as well as angiogenesis, may be an important population within EcE to sustain the growth and survival of endometriotic lesions. We observed altered metabolic activity and steroidogenesis predominantly in perivascular cells of EcE, supporting our hypothesis. These findings may facilitate further development of alternative strategies of non-hormonal treatment targeting metabolic pathways.

## Materials and methods

The study was approved by the Research Ethics Committee of the University of Tartu, Estonia (No 333/T-6), and written informed consent was obtained from all participants.

### Patient selection and sample processing

Both, the paired EuE and EcE samples were collected from four women with endometriosis undergoing laparoscopic surgery at the Tartu University Hospital (Tartu, Estonia). All women were in their follicular phase of the menstrual cycle (days 7 - 11), 33 ± 6.4 years old (mean ± standard deviation), with normal body mass index of 21 ± 1.8 kg/m2, and had not received any hormonal treatments for at least 3 months prior to the time of laparoscopy. According to the revised American Society for Reproductive Medicine classification system^55^, three women had minimal-mild endometriosis and one woman had moderate-severe endometriosis. During the laparoscopic surgery, peritoneal lesions were removed, and endometrial samples were collected using an endometrial suction catheter (Pipelle, Laboratoire CCD, France). All specimens were cut into two pieces and one portion was immediately fixed in 10% formalin for tissue histological evaluation, and the other portion stored in cryopreservation medium containing 1× Dulbecco’s Modified Eagle’s Medium (DMEM, Thermo Fisher Scientific), 30% fetal bovine serum (FBS, Biowest, Riverside, MO, USA) and 7.5% Dimethyl Sulfoxide Hybri-Max (Sigma-Aldrich). Tissue samples in cryopreservation medium were placed into Nalgene Cryo 1°C ‘Mr Frosty’ Freezing Container (Thermo Scientific) and deposited into a −80°C freezer overnight. The frozen biopsies were then stored in liquid nitrogen until further use. The presence of endometriosis-specific morphological features in the lesions was confirmed by a pathologist.

### Tissue dissociation for scRNA-seq

Cryopreserved tissues (50 – 80 mg) were warmed in a water bath at 37°C until thawed. Immediately after thawing, the biopsy content was washed twice in a 15 ml falcon tube with a 5 - 7 ml pre-warmed DMEM medium (phenol red free, charcoal-stripped FBS 10% + antibiotics/antifungal, Corning), to remove cryoprotectant and excess blood. The tissues were dissociated using an enzymatic cocktail of 2.5 mg/ml collagenase I (Sigma), 0.25 mg/ml DNase (AppliChem GmbH) and 10 mg/ml Dispase II (Gibco) in 10 ml of the DMEM medium. After shaking the tubes vigorously, further dissociation was continued for around 1 hour in an incubator at 37°C by placing the tubes in the horizontal position on a rotating shaker. The resulting suspension of the tissues was left unagitated for 1 - 2 min in a vertical position to settle the undigested tissue fragments by gravity. Any undigested tissue material was re-digested enzymatically in an incubator for another 15 min. The single-cell suspension was filtered through a 30-μm cell strainer to remove debris and cells were pelleted by centrifugation at 300×g for 5 min. Red blood cells were removed by the treatment with ACK lysis buffer (Gibco), following the manufacturer’s instructions. Pelleted single cells were then resuspended in 500 μl of DMEM medium and filtered again through a 30 μm cell strainer to collect the suspension in a 5% BSA-coated 1.5 ml microfuge tube. The cells were counted using Trypan blue chemistry on the automated counter (Corning) to determine the concentration and viability of the cell suspension. Live cells were enriched using MACS dead cell removal kit (Miltenyi Biotec) following the manufacturer’s instructions. The enriched cell suspensions having viability > 90% were washed with 0.04% BSA (Sigma) in Dulbecco’s DPBS (1×, Gibco) on ice for three times to remove ambient RNAs. The cells were resuspended in 0.04% BSA in DPBS to achieve a final concentration of 700 - 1200 cells/μl, as recommended by the manufacturer’s protocol (10x Genomics).

### Chromium 10x single-cell capturing, library generation and sequencing

Single cell RNA libraries were constructed using 10x Chromium Next GEM Single Cell 3’ reagent v3.1 kit (Dual index, 10x Genomics, CG000315 Rev C) comprised of single cell 3’ gel bead kit, library construction kit and chip G kit as per the manufacturer’s instructions (10x Genomics). Single-cell suspensions, single-cell 3’ gel beads, and master mix for reverse transcription reagents were loaded onto a 10x Chromium microfluidic chip for a targeted cell recovery of 3,000 cells per sample. The samples were processed in a 10x Chromium controller to generate Gel Beads-in-emulsion (GEM). Further steps, including cell lysis, first-strand cDNA synthesis, amplification, and purification, were carried out as per the manufacturer’s instructions to produce barcoded full-length cDNA. Library preparation was carried out simultaneously for all the samples using the library construction kit according to manufacturer’s instructions to avoid the batch effect. Quality control of cDNA and single-cell libraries were analyzed using Agilent 4150 tape station (Agilent Technologies). Dual indexed single-cell libraries were pooled, and pair-end sequenced targeting 35,000 reads per cell using NovaSeq PE150 (Illumina).

### Quality control, normalization, doublet cell removal, sample integration, and cell-type clustering

The 10x Genomics scRNA-seq data was aligned with refdata-gex-GRCh38-2020-A reference genome downloaded from the 10x website, and count matrices for each sample were created using the Cell Ranger (v7.0.0) software.

The 10x Genomics scRNA-seq data were analyzed with the CRAN package Seurat. In the data processing procedure, we retained cells with a UMI count of more than 500 and cells with the number of genes more than 200. High-complexity cell types (> 0.80) were kept, and cells with more than 15% of mitochondrial reads were filtered out. Genes with zero counts were removed by keeping only genes that were expressed within 3 or more cells. Following which, each sample was normalized, and the cell cycle score was assigned by the CellCycleScoring function based on the expression of G2/M and S phase markers collected from Tirosh et. al., according to the Seurat package manual^56^. Sctransform method of normalizing, estimating the variance of the raw filtered data, and identifying the most variable genes was applied to regress out the source of variation by mitochondrial expression, and cell cycle phase. Next, the Sctransformed samples were subjected to Principal Component Analysis (PCA), and a minimum number (n = 40) of PCs were identified. Based on the resulting PCs, we ran UMAP clustering, FindNeighbors, and FindClusters with 0.1 resolution steps from Seurat. Next, the pre-processed data was submitted to the DoubletFinder^57^ R package with default parameters. We then removed the identified cell doublets from each sample before integrating all eight samples. 5,000 highly variable shared features were used to integrate the samples to identify shared subpopulations across EuE and EcE. Canonical correlation analysis (CCA) was performed to identify shared sources of variation between the groups. Briefly, CCA identifies the greatest sources of variation in the data, but only if it is shared or conserved across the conditions/groups using the 5,000 variable features. In the second step of integration, the mutual nearest neighbors (MNN) were identified, and incorrect anchors (cells) were filtered out to integrate the samples across conditions. After integration, to visualize the integrated data, we used dimensionality reduction techniques, PCA, and UMAP. FindClusters with resolutions ranging from 0.2 to 1.4 from Seurat were used to find clusters in the integrated data. The UMAP technique was used to visualize the cell clusters. This integrated dataset with identified cell clusters was used for further downstream analysis.

### Cell cluster annotations

We used two approaches to identify and label the major cell types in the integrated dataset. First, the FindAllMarkers function from the Seurat package was used to find the differentially expressed genes from each identified cell class with default parameters. Next, we prepared a cell marker list collected from literature^6,7,27,58,59^, Cell Marker database^60^, Panglao database^61^, and single-cell RNA database from the Human protein atlas to validate the identified cell markers (Suppl. Table 6). To shortlist the marker genes, we applied log_2_FC > 1, adjusted p-value < 0.05, PCT.1 ≥ 0.7 (higher expression of a marker in a specific cluster) and PCT.2 ≤

0.3 (lower expression of the same marker in other clusters). Statistical analysis of cell proportions, comparison of total cell numbers and cell numbers by cell cycle phases between EcE and EuE was performed using Fisher’s exact test.

### Pseudobulk differential expression and KEGG pathway analyses

Pseudobulk differential expression analysis between the cell clusters of EcE and EuE and for each cell cycle phase was carried out using DESeq2 (v.1.40.1) R package^62^. The gene enrichment and the KEGG pathway enrichment analyses of differentially regulated genes (adjusted p-value < 0.05, log_2_FC ≤ –0.40 and ≥ 0.40) in each cell cluster were performed using clusterProfiler (v.4.8.1)^63,64^ R package.

### Metabolic and steroidogenesis pathways analysis

We selected 12 pathways related to cellular energy metabolism from the KEGG pathway database available online (Suppl. Table 7). For each pathway gene list, we extracted the differential expressions in EcE vs EuE from the output of DESeq2 pseudobulk differential expression analysis as mentioned in the previous step. Similarly, the genes involved in steroidogenesis were checked (Suppl. Table 8).

### The single-cell pathway analysis

We used the SCPA method^65^ to rank the 12 selected metabolic pathways between EcE and EuE, in each cell cluster. SCPA analysis assesses the changes in the multivariate distribution of a pathway and not just pathway enrichment. We used Q-value (multivariate distribution) measurement as a primary statistic, as suggested by the authors of the SCPA method.

## Data availability

Supplementary data is available online. The raw transcriptomic data was deposited at the GEO repository and is publicly available under the GEO accession number GSE247695.

## Acknowledgements

We acknowledge the use of BioRender.com to create the Figure 1A, Figure 2C and Figure 3. This research was supported by the European Union’s Horizon 2020 research and innovation

MATER program under grant agreement No. 813707 (M.Sar.), Estonian Research Council grants PRG1076 (A.S.), MOBJD1056 (A.P.), Horizon 2020 innovation grant ERIN, grant no. EU952516 (A.S.), Enterprise Estonia grant no EU48695 (M.P. and M.Saa.) and MSCA-RISE-2020 project TRENDO grant no 101008193 (A.S.).

## Contributions

A.S., M.P., M.Sar., and A.L. designed the study. P.S. collected samples and clinical data from patients, analysed clinical data. M.Sar., A.P., and M.Saa. performed wet-lab experiments. A.L., A.P. and V.M. performed the bioinformatic analysis. M.Sar., A.P., M.Saa., and M.P. performed the analysis and interpretation of bioinformatic results. M.Sar., A.L., A.P. and M.Saa. wrote the manuscript. M.Sar., A.L. and A.P. visualized the graphical data. M.P., M.Saa., A.S., K.G.D., P.G.L.L., T.K. and A.T. contributed to the interpretation of study results and discussions, revised and edited the manuscript. All authors read and approved the final version of the manuscript.

## Supplementary data

**Supplementary Table 1.** Total cell counts and percentages for all clusters in ectopic endometrium (EcE) and eutopic endometrium (EuE). Fisher’s exact test was used to compare cell cluster proportions between EcE and EuE.

**Supplementary Table 2.** Analysis of differential expression of 12 metabolic pathways between ectopic endometrium (EcE) and eutopic endometrium (EuE). Numbers of differentially expressed genes of metabolic pathways and percentages of the genes calculated to the total N of genes in a pathway.

**Supplementary Table 3.** SCPA analysis of the activity of 12 metabolic pathways in ectopic endometrium vs eutopic endometrium (by cell clusters).

**Supplementary Table 4.** Statistically significant differentially regulated genes of metabolic pathways between ectopic endometrium and eutopic endometrium. NS - statistically non-significant value (padj < 0.05).

**Supplementary Table 5.** Total cell counts and percentages for perivascular, stromal and endothelial clusters in ectopic endometrium (EcE) and eutopic endometrium (EuE) by cell cycle phases. Fisher’s exact test was used to compare cell cluster proportions between EcE and EuE for each cell cycle phase.

**Supplementary Table 6.** Gene markers used for cell cluster annotation. The cluster names are given as unmerged (upper line) and merged (bottom line).

**Supplementary Table 7.** A list of metabolic pathways obtained from KEGG pathway database, Homo sapiens (human).

**Supplementary Table 8.** A list of genes from steroidogenesis pathway.

**Supplementary Figure 1.** UMAP of 16 unmerged clusters. The clusters of Eutopic endometrium – on the left side, ectopic endometrium – on the right side.

**Supplementary Figure 2.** Differentially expressed gene markers in 16 unmerged clusters. Each dot shows the average expression and percent expressed. Each cluster exhibited the expression of cell-type representative marker genes.

**Supplementary Figure 3.** Top 10 enriched KEGG pathways in peritoneal, stromal and endothelial cell clusters of ectopic endometrium compared to eutopic endometrium.

**Supplementary Figure 4.** Venn diagram of top 10 enriched KEGG pathways in perivascular, stromal and endothelial cell clusters of ectopic endometrium compared to eutopic endometrium.

**Supplementary Figure 5.** Top 20 DEGs of perivascular, stromal and endothelial clusters. The clusters of Eutopic endometrium (EuE) – in red, ectopic endometrium (EcE) – in blue.

## References

1. Zondervan, K. T., Becker, C. M. & Missmer, S. A. Endometriosis. N Engl J Med 382, 1244–1256 (2020).

2. Giudice, L. C. & Kao, L. C. Endometriosis. The Lancet 364, 1789–1799 (2004).

3. Saunders, P. T. K. & Horne, A. W. Endometriosis: Etiology, pathobiology, and therapeutic prospects. Cell 184, 2807–2824 (2021).

4. Fonseca, M. A. S. et al. Single-cell transcriptomic analysis of endometriosis. Nat Genet 55, 255–267 (2023).

5. Queckbörner, S. et al. Stromal Heterogeneity in the Human Proliferative Endometrium— A Single-Cell RNA Sequencing Study. JPM 11, 448 (2021).

6. Tan, Y. et al. Single-cell analysis of endometriosis reveals a coordinated transcriptional programme driving immunotolerance and angiogenesis across eutopic and ectopic tissues. Nat Cell Biol 24, 1306–1318 (2022).

7. Wang, W. et al. Single-cell transcriptomic atlas of the human endometrium during the menstrual cycle. Nat Med 26, 1644–1653 (2020).

8. Jabbour, H. N., Kelly, R. W., Fraser, H. M. & Critchley, H. O. D. Endocrine Regulation of Menstruation. Endocrine Reviews 27, 17–46 (2006).

9. Zhu, J. & Thompson, C. B. Metabolic regulation of cell growth and proliferation. Nat Rev Mol Cell Biol 20, 436–450 (2019).

10. Hsu, C.-Y. et al. Synthetic Steroid Hormones Regulated Cell Proliferation Through MicroRNA-34a-5p in Human Ovarian Endometrioma. Biol Reprod 94, 60 (2016).

11. Saha, S., Dey, S. & Nath, S. Steroid Hormone Receptors: Links With Cell Cycle Machinery and Breast Cancer Progression. Front Oncol 11, 620214 (2021).

12. Weigel, N. L. & Moore, N. L. Cyclins, cyclin dependent kinases, and regulation of steroid receptor action. Mol Cell Endocrinol 265–266, 157–161 (2007).

13. Fajas-Coll, L., Lagarrigue, S., Hure, S., Lopez-Mejía, I. & Denechaud, P.-D. Cell Cycle and Metabolic Changes During Tissue Regeneration and Remodeling. in Pathobiology of Human Disease (eds. McManus, L. M. & Mitchell, R. N.) 542–549 (Academic Press, San Diego, 2014). doi:10.1016/B978-0-12-386456-7.02101-8.

14. Kalucka, J. et al. Metabolic control of the cell cycle. Cell Cycle 14, 3379–3388 (2015).

15. Critchley, H. O. D., Maybin, J. A., Armstrong, G. M. & Williams, A. R. W. Physiology of the Endometrium and Regulation of Menstruation. Physiological Reviews 100, 1149– 1179 (2020).

16. Rižner, T. L. Estrogen metabolism and action in endometriosis. Molecular and Cellular Endocrinology 307, 8–18 (2009).

17. Tsai, S. J., Wu, M. H., Lin, C. C., Sun, H. S. & Chen, H. M. Regulation of steroidogenic acute regulatory protein expression and progesterone production in endometriotic stromal cells. J Clin Endocrinol Metab 86, 5765–5773 (2001).

18. Wu, M.-H., Hsiao, K.-Y. & Tsai, S.-J. Hypoxia: The force of endometriosis. J Obstet Gynaecol Res 45, 532–541 (2019).

19. Lee, H.-C., Lin, S.-C., Wu, M.-H. & Tsai, S.-J. Induction of Pyruvate Dehydrogenase Kinase 1 by Hypoxia Alters Cellular Metabolism and Inhibits Apoptosis in Endometriotic Stromal Cells. Reprod Sci 26, 734–744 (2019).

20. Kasvandik, S. et al. Deep Quantitative Proteomics Reveals Extensive Metabolic Reprogramming and Cancer-Like Changes of Ectopic Endometriotic Stromal Cells. J Proteome Res 15, 572–584 (2016).

21. Young, V. J. et al. Transforming Growth Factor-β Induced Warburg-Like Metabolic Reprogramming May Underpin the Development of Peritoneal Endometriosis. J Clin Endocrinol Metab 99, 3450–3459 (2014).

22. Vander Heiden, M. G., Cantley, L. C. & Thompson, C. B. Understanding the Warburg Effect: The Metabolic Requirements of Cell Proliferation. Science 324, 1029–1033 (2009).

23. Vaupel, P., Schmidberger, H. & Mayer, A. The Warburg effect: essential part of metabolic reprogramming and central contributor to cancer progression. Int J Radiat Biol 95, 912–919 (2019).

24. Horne, A. W. et al. Repurposing dichloroacetate for the treatment of women with endometriosis. Proc Natl Acad Sci U S A 116, 25389–25391 (2019).

25. Kobayashi, H., Shigetomi, H. & Imanaka, S. Nonhormonal therapy for endometriosis based on energy metabolism regulation. Reproduction and Fertility 2, C42–C57 (2021).

26. McKinnon, B. et al. Glucose transporter expression in eutopic endometrial tissue and ectopic endometriotic lesions. J Mol Endocrinol 52, 169–179 (2014).

27. Ma, J. et al. Single-cell transcriptomic analysis of endometriosis provides insights into fibroblast fates and immune cell heterogeneity. Cell & Bioscience 11, 125 (2021).

28. Zhu, S. et al. The heterogeneity of fibrosis and angiogenesis in endometriosis revealed by single-cell RNA-sequencing. Biochim Biophys Acta Mol Basis Dis 1869, 166602 (2023).

29. Avolio, E., Alvino, V. V., Ghorbel, M. T. & Campagnolo, P. Perivascular cells and tissue engineering: Current applications and untapped potential. Pharmacol Ther 171, 83–92 (2017).

30. Masuda, H., Anwar, S. S., Bühring, H.-J., Rao, J. R. & Gargett, C. E. A novel marker of human endometrial mesenchymal stem-like cells. Cell Transplant 21, 2201–2214 (2012).

31. Cousins, F. L., O, D. F. & Gargett, C. E. Endometrial stem/progenitor cells and their role in the pathogenesis of endometriosis. Best Practice & Research Clinical Obstetrics & Gynaecology 50, 27–38 (2018).

32. Cousins, F. L., Filby, C. E. & Gargett, C. E. Endometrial Stem/Progenitor Cells–Their Role in Endometrial Repair and Regeneration. Front Reprod Health 3, 811537 (2022).

33. Jones, R. G. et al. AMP-Activated Protein Kinase Induces a p53-Dependent Metabolic Checkpoint. Molecular Cell 18, 283–293 (2005).

34. Herzig, S. & Shaw, R. J. AMPK: guardian of metabolism and mitochondrial homeostasis. Nat Rev Mol Cell Biol 19, 121–135 (2018).

35. Assaf, L., Eid, A. A. & Nassif, J. Role of AMPK/mTOR, mitochondria, and ROS in the pathogenesis of endometriosis. Life Sciences 306, 120805 (2022).

36. Chandel, N. S. Evolution of Mitochondria as Signaling Organelles. Cell Metab 22, 204– 206 (2015).

37. Young, V. J. et al. Transforming Growth Factor-β Induced Warburg-Like Metabolic Reprogramming May Underpin the Development of Peritoneal Endometriosis. J Clin Endocrinol Metab 99, 3450–3459 (2014).

38. Atkins, H. M. et al. Endometrium and endometriosis tissue mitochondrial energy metabolism in a nonhuman primate model. Reproductive Biology and Endocrinology 17, 70 (2019).

39. Yu, L. et al. Modeling the Genetic Regulation of Cancer Metabolism: Interplay Between Glycolysis and Oxidative Phosphorylation. Cancer Res 77, 1564–1574 (2017).

40. Xiao, Z., Dai, Z. & Locasale, J. W. Metabolic landscape of the tumor microenvironment at single cell resolution. Nat Commun 10, 3763 (2019).

41. Cai, X. et al. Lactate activates the mitochondrial electron transport chain independently of its metabolism. Molecular Cell 83, 3904–3920.e7 (2023).

42. Zeitoun, K. et al. Deficient 17β-Hydroxysteroid Dehydrogenase Type 2 Expression in Endometriosis: Failure to Metabolize 17β-Estradiol1. The Journal of Clinical Endocrinology & Metabolism 83, 4474–4480 (1998).

43. Kyama, C. M. et al. Endometrial and peritoneal expression of aromatase, cytokines, and adhesion factors in women with endometriosis. Fertil Steril 89, 301–310 (2008).

44. Delvoux, B. et al. Increased Production of 17β-Estradiol in Endometriosis Lesions Is the Result of Impaired Metabolism. The Journal of Clinical Endocrinology & Metabolism 94, 876–883 (2009).

45. Bulun, S. E. et al. Estrogen Receptor-β, Estrogen Receptor-α, and Progesterone Resistance in Endometriosis. Semin Reprod Med 28, 36–43 (2010).

46. Kazi, A. A., Molitoris, K. H. & Koos, R. D. Estrogen Rapidly Activates the PI3K/AKT Pathway and Hypoxia-Inducible Factor 1 and Induces Vascular Endothelial Growth Factor A Expression in Luminal Epithelial Cells of the Rat Uterus. Biol Reprod 81, 378– 387 (2009).

47. Monsivais, D. et al. ERβ- and Prostaglandin E2-Regulated Pathways Integrate Cell Proliferation via Ras-like and Estrogen-Regulated Growth Inhibitor in Endometriosis. Mol Endocrinol 28, 1304–1315 (2014).

48. Zhang, X. et al. 17β-Estradiol promotes angiogenesis of bone marrow mesenchymal stem cells by upregulating the PI3K-Akt signaling pathway. Computational and Structural Biotechnology Journal 20, 3864–3873 (2022).

49. Maekawa, R. et al. Aberrant DNA methylation suppresses expression of estrogen receptor 1 (ESR1) in ovarian endometrioma. Journal of Ovarian Research 12, 14 (2019).

50. Yilmaz, B. D. & Bulun, S. E. Endometriosis and nuclear receptors. Human Reproduction Update 25, 473–485 (2019).

51. Lv, H. et al. Deciphering the endometrial niche of human thin endometrium at single-cell resolution. Proceedings of the National Academy of Sciences 119, e2115912119 (2022).

52. McKinnon, B. D. et al. Altered differentiation of endometrial mesenchymal stromal fibroblasts is associated with endometriosis susceptibility. Commun Biol 5, 1–14 (2022).

53. Mehrmohamadi, M., Liu, X., Shestov, A. A. & Locasale, J. W. Characterization of the usage of the serine metabolic network in human cancer. Cell Rep 9, 1507–1519 (2014).

54. Peng, X. et al. Molecular Characterization and Clinical Relevance of Metabolic Expression Subtypes in Human Cancers. Cell Rep 23, 255–269.e4 (2018).

55. Revised American Society for Reproductive Medicine classification of endometriosis: 1996. Fertil Steril 67, 817–821 (1997).

56. Tirosh, I. et al. Dissecting the multicellular ecosystem of metastatic melanoma by single-cell RNA-seq. Science 352, 189–196 (2016).

57. McGinnis, C. S., Murrow, L. M. & Gartner, Z. J. DoubletFinder: Doublet Detection in Single-Cell RNA Sequencing Data Using Artificial Nearest Neighbors. Cell Syst 8, 329–337.e4 (2019).

58. Garcia-Alonso, L. et al. Mapping the temporal and spatial dynamics of the human endometrium in vivo and in vitro. Nat Genet 53, 1698–1711 (2021).

59. Zou, G. et al. Cell subtypes and immune dysfunction in peritoneal fluid of endometriosis revealed by single-cell RNA-sequencing. Cell & Bioscience 11, 98 (2021).

60. Hu, C. et al. CellMarker 2.0: an updated database of manually curated cell markers in human/mouse and web tools based on scRNA-seq data. Nucleic Acids Res 51, D870– D876 (2023).

61. Franzén, O., Gan, L.-M. & Björkegren, J. L. M. PanglaoDB: a web server for exploration of mouse and human single-cell RNA sequencing data. Database 2019, baz046 (2019).

62. Love, M. I., Huber, W. & Anders, S. Moderated estimation of fold change and dispersion for RNA-seq data with DESeq2. Genome Biol 15, 550 (2014).

63. Yu, G., Wang, L.-G., Han, Y. & He, Q.-Y. clusterProfiler: an R package for comparing biological themes among gene clusters. OMICS 16, 284–287 (2012).

64. Wu, T. et al. clusterProfiler 4.0: A universal enrichment tool for interpreting omics data. Innovation (Camb) 2, 100141 (2021).

65. Bibby, J. A., et al. Systematic single-cell pathway analysis to characterize early T cell activation. Cell Reports 41, (2022).

