## Supplementary figures and images for "Endometriotic lesions exhibit distinct metabolic signature compared to paired eutopic endometrium at the single-cell level"

### Supp. Fig. 1

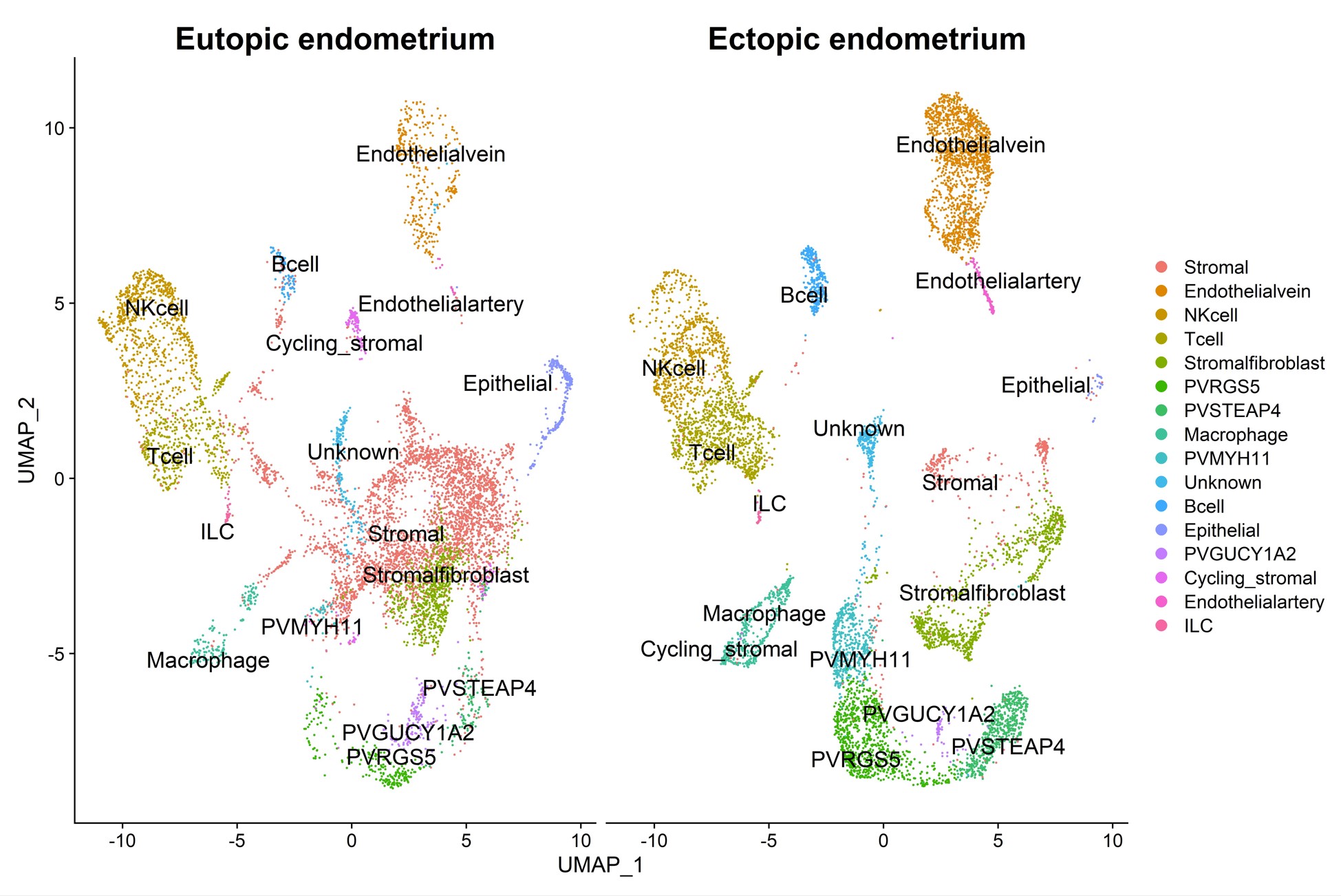

### Supp. Fig. 2

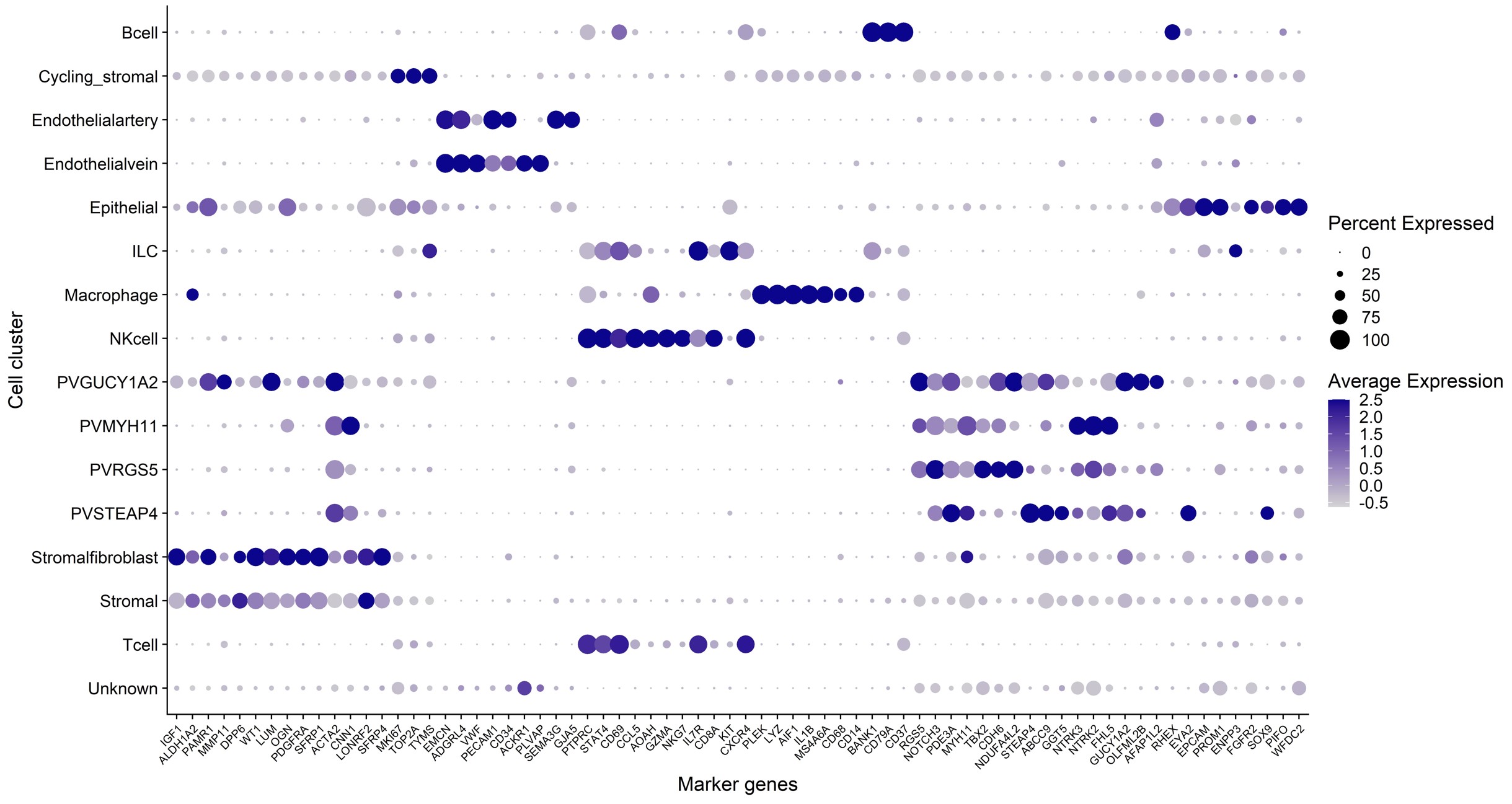

### Supp. Fig. 3

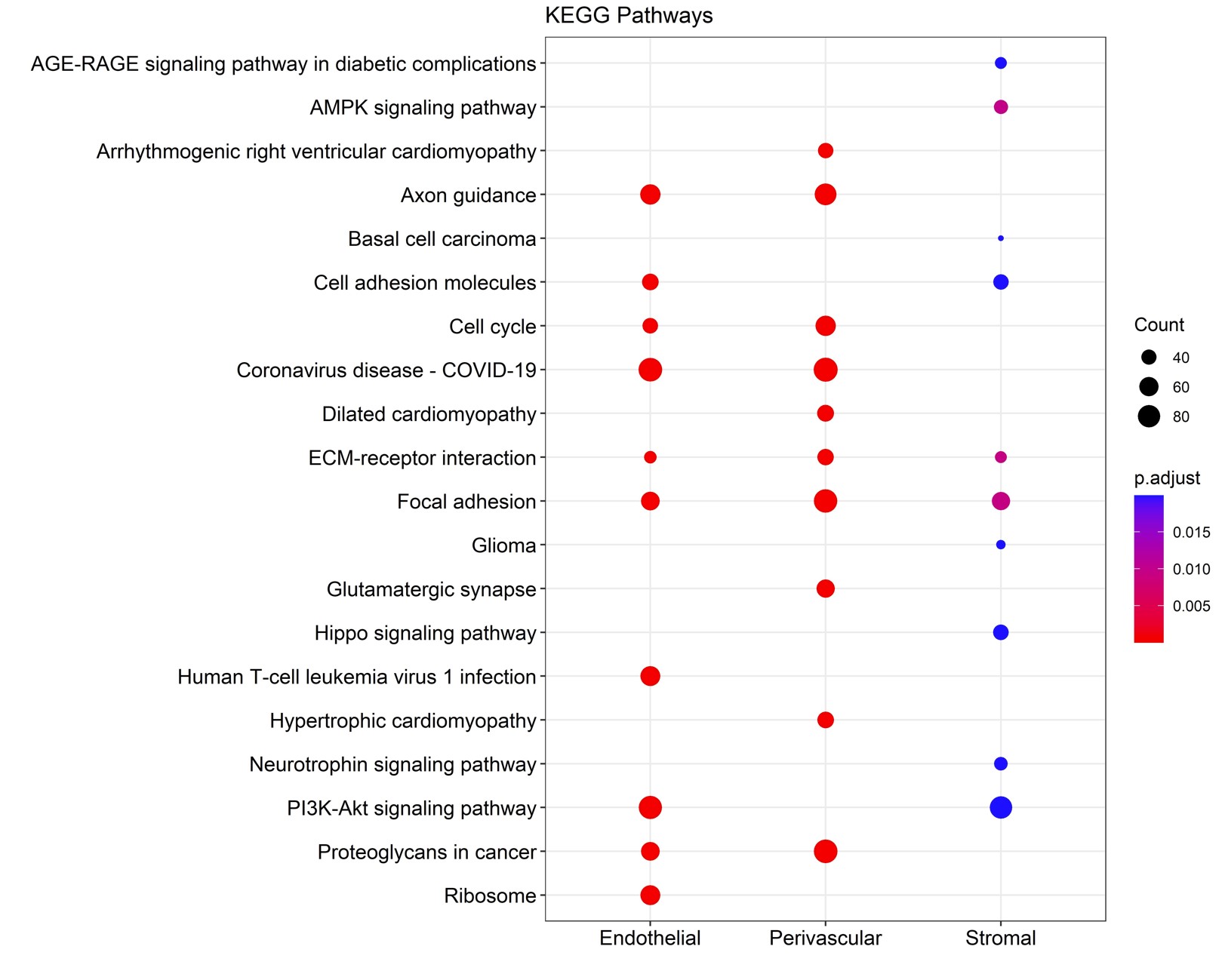

### Supp. Fig. 4

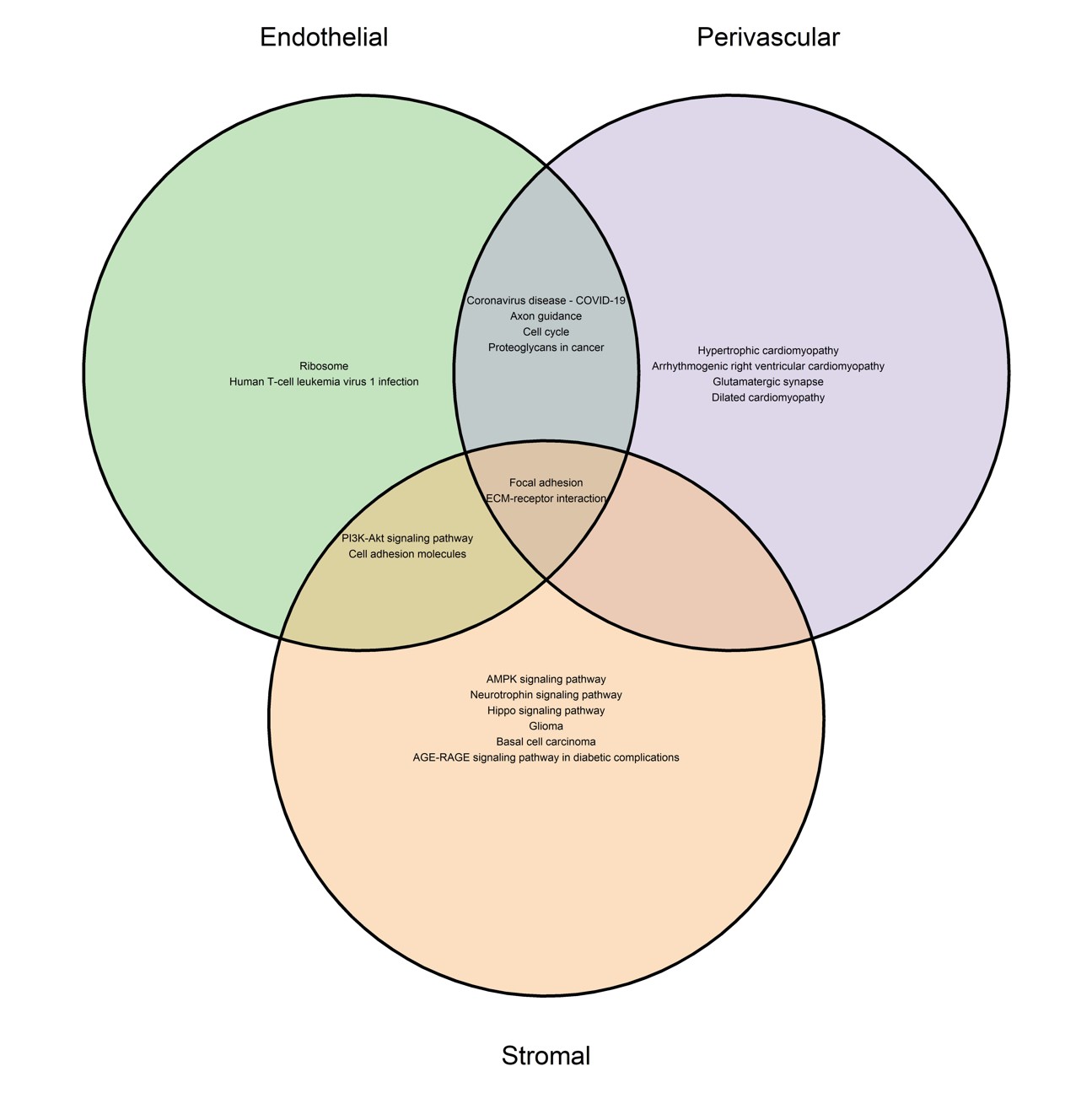

### Supp. Fig. 5

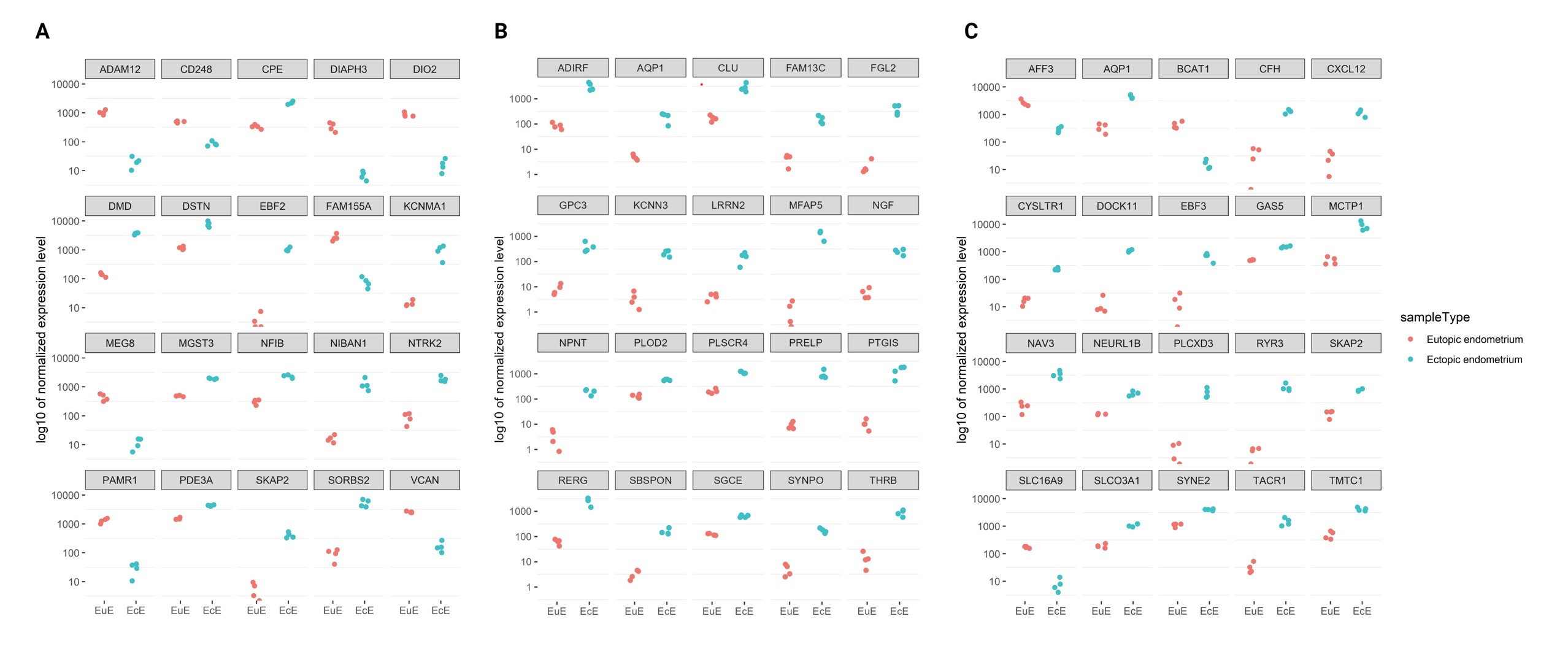
